# Lipid Hydrogen Stable Isotope Probing Reveals Archaeal Lipid Turnover in Hot Spring Sediments

**DOI:** 10.64898/2026.05.15.725266

**Authors:** Carolynn M. Harris, Sebastian Kopf, Maximiliano J. Amenabar, Eric S. Boyd, Xiahong Feng, Ann Pearson, William D. Leavitt

## Abstract

Quantifying the lipid biosynthesis rate of archaea in hot spring sediments is necessary to estimate their generation times and environmental significance, with implications for modern biogeochemical cycling and astrobiology. Here, we performed lipid hydrogen stable isotope probing (LH-SIP) experiments on sediments collected from two high-temperature, suboxic, circumneutral hot springs in Yellowstone National Park (USA) and El Tatio Geyserfield (Chile). We determined the incorporation rate of ^2^H_2_O into intact polar lipids (IPLs) during short-duration, *ex situ* incubations, which provides a taxon-and metabolism-agnostic quantification of biosynthesis under near-natural conditions. We targeted isoprenoid glycerol dialkyl glycerol tetraether lipids (iGDGTs) and recovered structures with 0 to 7 cyclopentyl moieties from both springs. We observed minor ^2^H-uptake into archaeal IPL-iGDGT-derived biphytanes in spring sediments in Yellowstone, corresponding to decadal-scale apparent generation times (16 ± 3 years), and no detectable uptake in El Tatio sediments, where lipid turnover was slower than could be resolved under the experimental conditions (maximum resolvable apparent generation times ∼ 42 ± 21 years). We conclude that these slow apparent generation times likely reflect substantial dilution by a large standing IPL pool and/or recycling or re-functionalization of relict lipid components, rather than equivalently slow rates of archaeal cell division. These are the first estimates of archaeal lipid production rates in terrestrial hydrothermal systems using LH-SIP incubations and provide critical constraints for interpreting archaeal lipids in modern systems and ancient hot spring deposits.

**Plain Language Summary:** Hot springs on Earth are important natural laboratories for understanding how signs of life might form and be preserved in hydrothermal environments on early Earth or Mars. In this study, we examine the rate of archaeal lipid production in sediments from two hot springs in Yellowstone National Park and the El Tatio Geyserfield in Chile. We used a method that measures new microbial lipid production by tracing heavy hydrogen from labeled water into newly made lipids in cell membranes. We found that archaeal lipids in hot spring sediments have very slow apparent turnover, on timescales of decades. These apparent turnover times likely reflect not only slow lipid synthesis, but also dilution by a large standing pool of older lipids and recycling of relict lipid components. This result, along with the chemical stability of lipids and the rapid mineralization rate in some hot springs, may allow these molecular biosignatures to be entombed and preserved in hot spring mineral deposits. These results help us constrain interpretations of ancient hydrothermal deposits on Earth and support the idea that microbial communities with slow rates of new lipid production could still leave detectable molecular traces in similar environments on Mars and other rocky planets.

**Key Points:**

- Lipid hydrogen stable isotope probing is applied to high temperature hot spring sediments for the first time
- Sedimentary archaeal lipids in hot springs exhibit apparent turnover on decadal timescales comparable to some marine sediments
- Terrestrial hot spring lipids provide a framework for evaluating potential preserved biosignatures in ancient Martian hydrothermal systems

## 1 Introduction

Terrestrial (subaerial) hot springs host diverse microbial communities that are adapted to multiple environmental extremes (Des Marais and Walter, 2019; Sriaporn et al., 2023). These communities of Bacteria and Archaea must survive in temperatures near the boiling point of water, acidic or alkaline pH, strongly oxidizing or reducing conditions, and often heavy-metal concentrations (Cady et al., 2018; Des Marais and Walter, 2019; Merino et al., 2019). Energy availability can be extremely limiting in such environments (Shock et al., 2010), and Archaea appear to be particularly well-adapted to these constraints (Valentine, 2007). The growth rate of these archaea is a fundamental ecological parameter critical for interpreting microbial contributions to carbon and energy fluxes in hot spring systems, including the production of organic biomarkers. Determining archaeal growth rates in hot springs is challenging because most inhabitants are uncultured and have relatively low abundance in sediments (Des Marais and Walter, 2019) due to the multiple physicochemical extremes they experience – including high temperature, low or high pH, low organic carbon, and suboxic to anoxic conditions (Cady et al., 2018; Pirajno, 2020).

Hot springs are useful analog environments for early Earth and hydrothermal systems beyond Earth, including fossil systems on Mars, that may host or have hosted habitable conditions (Pirajno, 2020; Finkel et al., 2023; Rogers et al., 2023; Beck et al., 2025). If hydrothermal systems beyond Earth once harbored life, it may have resembled the thermophilic archaea in modern hot springs (Siliakus et al., 2017; Matthews et al., 2023). Even slow-growing archaea may produce faint or cryptic molecular signals that could persist in the geological record. Better estimates of archaeal growth rates in hot spring sediments will allow us to refine both biosignature detection thresholds and an interpretive framework for planetary missions. Accurate growth rate determinations from extant archaea across springs with diverse geochemistry will help characterize what conditions are sufficient to support observable life, how rapidly organic biosignatures form, and how much biomass accumulates and is preserved in mineral deposits (e.g., sinters) under conditions relevant to early Earth and Martian hydrothermal systems.

In active hot springs, high temperature (typically >70 °C) precludes colonization by phototrophs (Cox et al., 2011; Boyd et al., 2012; Fecteau et al., 2022), and pH, temperature, and energy availability are the dominant parameters controlling the structure and composition of chemotrophic microbial communities (Swingley et al., 2012; Inskeep et al., 2013; Colman et al., 2024; Qi et al., 2024). Hot springs often host relatively simple communities in which archaea are prominent or dominant, including Thermoproteota (e.g., Sulfolobales) in high-temperature acidic to circumneutral springs, methanogenic Euryarchaeota in anoxic springs, and ammonia-oxidizing Thaumarchaeota in cooler, oxic springs (Mueller et al., 2021; Sriaporn et al., 2023).

Many of these archaea can synthesize a characteristic series of membrane lipids called isoprenoid glycerol dialkyl glycerol tetraethers (iGDGTs), comprising two C_40_ biphytanyl (BP) chains (Schouten et al., 2007; Podar et al., 2020; Chong, 2024). Common iGDGTs contain from 0 to 8 cyclopentane moieties (i.e., iGDGT-0 to iGDGT-8) with up to 4 rings per biphytane (i.e., BP-0 to BP-4) (Schouten et al., 2013). In response to heat or acid stress, cells regulate the fluidity, permeability, and rigidity of their cell membranes by synthesizing more highly ringed structures (Boyd et al., 2011; Cobban et al., 2020; Zhou et al., 2020), which physically pack tighter (Gabriel and Chong, 2000; Gliozzi et al., 1983; Zhao et al., 2025). Crenarchaeol is a structurally distinct iGDGT, containing four cyclopentane and one cyclohexane moiety, that is also common in hot springs, particularly those with lower temperature and circumneutral pH (Pearson and Ingalls, 2013; Schouten et al., 2013; Calhoun et al., 2026). Thus, the distribution of iGDGT structures in hot springs is related to temperature, pH, and other stressors (Pearson et al., 2008; Boyd et al., 2013; Kaur et al., 2015; Hurley et al., 2016; Elling et al., 2017). iGDGT lipids are also robust to diagenesis, and in certain preservational settings can survive in the geologic record for hundreds of millions of years (Vinnichenko et al., 2020). As such, iGDGTs have utility as both microbial biosignatures and paleo-environmental proxies, offering insights into the presence, metabolism, and ecology of Archaea in modern and ancient environments (Pearson and Ingalls, 2013).

Hot spring mineral deposits – including siliceous sinters (geyserite), calcareous sinters (travertine), and iron oxides – can entomb and preserve these lipids (Hays et al., 2017; Cady et al., 2018; Des Marais and Walter, 2019). Indeed, microbes may contribute to the morphogenesis of mineral deposits proximal to spring vents (Campbell et al., 2015; Teece et al., 2023). Comparable spring deposits are likely to occur on other terrestrial planets, and several promising candidates have been identified on Mars (Ruff et al., 2020; Rogers et al., 2023). To interpret archaeal lipid biosignatures in modern sediments or ancient deposits requires better constraints on the rates of new production and persistence of individual lipid moieties, including tetraethers with different ring distributions. Production and preservation dynamics influence the abundance, isotopic composition, and environmental significance of lipid biosignatures on Earth and in extraterrestrial hydrothermal settings.

Direct measurements of archaeal growth rates in hot spring environments are rare, and most published growth rates pertain to planktonic cells or filaments in spring water columns, biofilm-associated cells in flowing water, or enrichment cultures in the lab (Shivvers and Brock, 1973; Bohlool and Brock, 1974; Mosser et al., 1974; Dunckel et al., 2009; Jay et al., 2015). While lab culturing is the most direct way to measure growth (yielding precise doubling times), these techniques only capture a subset of the archaeal community that grows rapidly on culture media often under optimal conditions and are not representative of natural growth rates in sediments (Figure S1). Many hot spring archaea remain uncultured even though they often account for the majority of biomass and biological activity in sediments (Colman et al., 2024). Previous studies have speculated that springs with high temperature and acidic pH may favor enhanced synthesis of iGDGTs over other membrane lipids, based on abundance patterns of core and intact polar lipids in spring sediments (Boyd et al., 2013) and in pure cultures (Fehyl-Buska, 2016). To date, there are no published studies providing direct quantitative estimates of archaeal growth rates in hot spring sediments. This knowledge gap is largely due to methodological constraints that prevent attribution of specific measurements to individual cells or populations in natural sediment communities.

Stable isotope probing (SIP) has emerged as a powerful approach to measure microbial growth and substrate utilization *in situ* or in *ex situ* microcosms without requiring isolation. In this approach, microbes are fed ^13^C, ^2^H, ^15^N, or ^18^O-labeled compounds (e.g., ^13^C-bicarbonate, ^13^C-organic substrates, ^2^H-water, etc.) and the incorporation of the heavy isotope into biomass is measured over time. SIP methods provide estimates of biosynthesis or metabolic turnover that are culture-independent and independent of the population dynamics of the target communities (i.e., steady state vs. growth vs. decline). Recent advances in SIP methodology couple traditional **l**ipid biomarker analysis to **<u>h</u>**ydrogen **<u>SIP</u>** (e.g., LH-SIP) to measure growth rates as incorporation of deuterated water (^2^H_2_O) (Kopf et al., 2015; Wegener et al., 2016; Caro et al., 2023). LH-SIP is agnostic to both metabolism and taxa because all known biomolecules incorporate hydrogen and no additional carbon substrates are provided. This technique is especially useful for uncultured populations and for observing growth under near-natural conditions. LH-SIP has successfully detected microbial activity in environments with low biomass and slow growth rates including alpine tundra (generation time ∼50 days) (Caro et al., 2023), hydrothermally influenced marine sediments (380 days to 28 years) (Kellermann et al., 2016), and the deep subsurface (160 to 840 years) (Wegener et al., 2012).

A handful of studies have applied SIP with various compounds to interrogate the activity of archaea in hydrothermal environments. For example, Schubotz and others amended mixed archaeal-bacterial “streamer” biofilm communities from Octopus Spring and ‘Bison Pool’ (both ∼80 °C, alkaline, silicious hot springs in Yellowstone), with various ^13^C-labeled substrates (bicarbonate, acetate, formate, glucose) (Schubotz et al., 2015). Assimilation of ^13^C-acetate into archaeal membrane lipids indicated growth via heterotrophic metabolisms in both springs (Schubotz et al., 2015). Researchers found widespread assimilation of ^13^C-labeled compounds into archaeal DNA in 74 °C sediments in Gongxiaoshe Hot Spring (China) (Lai et al., 2023). In another study, ^2^H_2_O SIP combined with single-cell sorting (via BONCAT-FACS, which fluorescently labels new proteins) identified active archaeal cells in a hot spring sediment community in the Five Sisters hot spring group in Yellowstone (72 °C, pH 8.5) (Reichart et al., 2020). Finally, metabolism of bicarbonate, formate, and/or acetate has been measured in a variety of high temperature hot spring communities, many of which were dominated by Archaea (Boyd et al., 2009; Urschel et al., 2015; Keller et al., 2023; Fernandes-Martins et al., 2023). While these studies reveal Archaea are active in hot springs, they do not provide direct estimates of generation times. Thus, the growth rate of archaea in sediments represents a key gap in our understanding of archaeal ecology in hot springs that has important implications for modern biogeochemical cycling and applications to astrobiology and life detection.

In this paper, we target two high-temperature (>80 °C), circumneutral (pH 6.5 to 7.1), suboxic (< 0.7 mg L^-1^ O_2_) hot springs in Yellowstone National Park (USA) and El Tatio Geyserfield (Chile) (Table 1, Figure 1). These springs provide an opportunity to examine lipid biosignature production by polyextremophilic archaea, which are promising candidates of life forms that could evolve beyond Earth (DasSarma et al., 2020; Matthews et al., 2023). We describe the geochemistry and archaeal lipid inventory in spring sediments and report the results of LH-SIP *ex situ* sediment incubations with ^2^H_2_O which were designed to provide compound-specific and community-integrated constraints on IPL-iGDGT biosynthesis and apparent generation or turnover times. We compare these apparent turnover times with those reported for archaeal lipids in other environments and discuss the processes that may decouple IPL-iGDGT lipid turnover from cellular growth in hot spring sediments.

**Figure 1.**
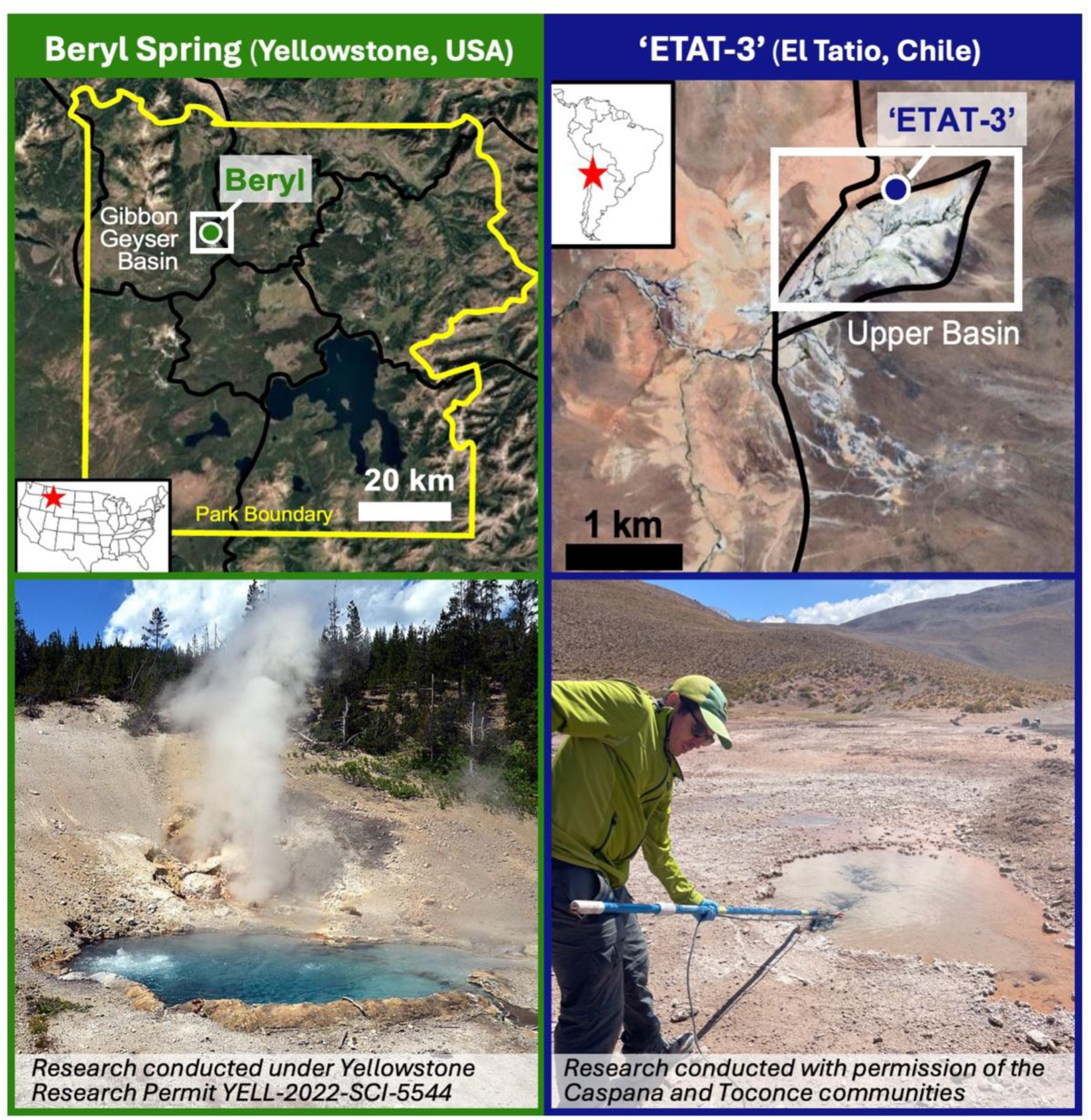
Field sites investigated in this study, including Beryl Spring in Yellowstone National Park, USA, and ‘ETAT-3’ in the El Tatio Geyserfield, Chile. Research in Yellowstone was conducted under Yellowstone Research Permit YELL-2022-SCI-5544; research in the El Tatio Geyserfield was conducted with permission of the Caspana and Toconce Communities. Beryl photo credited to Wikipedia. ‘ETAT-3’ photo credited to C. Harris.

**Table 1.** Site characteristics, aqueous geochemistry, and archaeal lipid inventory for the two hot springs examined in this study: Beryl Spring (Yellowstone National Park, USA) and ‘ETAT-3’ (El Tatio Geyserfield, Chile). ORP = oxidation/reduction potential; DO = dissolved oxygen; SpC = specific conductivity. Total isoprenoid glycerol dibiphytanyl glycerol tetraether (iGDGT) lipid abundance is reported as µg g^-1^ sediment, and IPL-iGDGT (%) represents the percentage of intact polar lipid iGDGTs relative to total iGDGTs. Apparent lipid–water hydrogen isotope fractionation (^2^ε_L/W_, ‰) between IPL-iGDGT-derived biphytanes and spring water measured at natural abundance is also reported. Values are reported as mean ± standard error where applicable.

| Parameter | Beryl | ‘ETAT-3’ |
| --- | --- | --- |
| Latitude (DD) | 44.679 | -22.328 |
| Longitude (DD) | -110.747 | -68.006 |
| Elevation (mASL) | 2239 | 4274 |
| T (°C) | 89.6 ± 0.1 | 79.7 ± 0.7 |
| pH | 6.5 ± 0.1 | 7.1 ± 0.03 |
| ORP (mV) | -334 ± 1 | -84 ± 9 |
| DO (mg L<sup>-1</sup>) | 0.4 ± 0.01 | 0.7 ± 0.02 |
| SpC (mS cm<sup>-1</sup>) | 2.5 ± 0.1 | 23.5 ± 0.2 |
| Water $\delta^2\text{H}$ (‰) | -139 ± 0.3 | -68 ± 0.1 |
| Water $\delta^{18}\text{O}$ (‰) | -14.6 ± 0.5 | -4.8 ± 0.5 |
| iGDGT lipids ($\mu\text{g g}^{-1}$ sed) | 80 | 12 |
| IPL-iGDGT (%) | 15% | 88% |
| $^2\epsilon_{\text{LW}}$ (‰) | -309 ± 3 | -255 ± 4 |

## 2 Methods

### 2.1 Sediment sampling

Sediment samples were collected from two hot springs: Beryl Spring (44.679 °N; -110.747 °W) in Yellowstone National Park (Wyoming, USA) in October 2022 and ‘ETAT-3’ hot spring (22.328 °S; -68.006 °W) in the El Tatio Geyserfield (Chile) in January 2023 (Table 1, Figure 1). These springs were selected because their high temperatures (>80 °C) preclude colonization by phototrophic microbes and favor thermophilic chemotrophic communities in which iGDGT-producing archaea are expected to be important contributors. At each spring, a dipstick with sterilized beaker was used to sample near-surface sediments from the margins of the active hot spring vent. Samples of the overlying spring water were collected in the same manner. Samples were stored in solvent-washed glass jars with Teflon-lined lids (DWK Life Sciences, PTFE liner) and placed in a cool, opaque insulated container before shipping to Dartmouth College, Hanover, NH. Approximately 3 days elapsed between collection and arrival at Dartmouth for the Yellowstone samples and approximately 7 days for the El Tatio samples. LH-SIP incubations were initiated within 24 hours of arrival.

A multiprobe data sonde (YSI, DSS Pro model) was used to determine major aqueous geochemistry parameters (temperature, pH, specific conductivity, dissolved oxygen, and redox potential) in spring water above the site of sediment sampling.

We also sampled a second Yellowstone spring, ‘Pentagonal’ (pH 7.8; 77.4 °C; 44.463°N; - 110.853°W), but recovered iGDGT abundances were insufficient for reliable structural and δ^2^H isotopic analyses, so those data are not discussed further.

### 2.2 Lipid hydrogen SIP incubations with ^2^H_2_O

The LH-SIP incubations were started within 24 hours of sample arrival at Dartmouth College following the approach of Caro et al., 2023, modified for anoxic conditions (Figure 2). In a Coy two-person anoxic chamber (95:5 N_2_:H_2_), sediments were sieved to 2 mm to remove large rocks, homogenized, weighed, and transferred to 70 mL glass serum bottles. Sediment and spring water were added at a 2:1 ratio, with 10 mL of filter-sterilized spring water per incubation and approximately 45 mL of headspace remaining in each bottle. The spring water was amended with 99.9% ^2^H_2_O to a nominal enrichment of 5,000 ppm ^2^H (0.5 atom % ^2^H; ∼31,000 ‰ VSMOW), although the final enrichment was lower due to dilution by sediment pore water.

**Figure 2.**
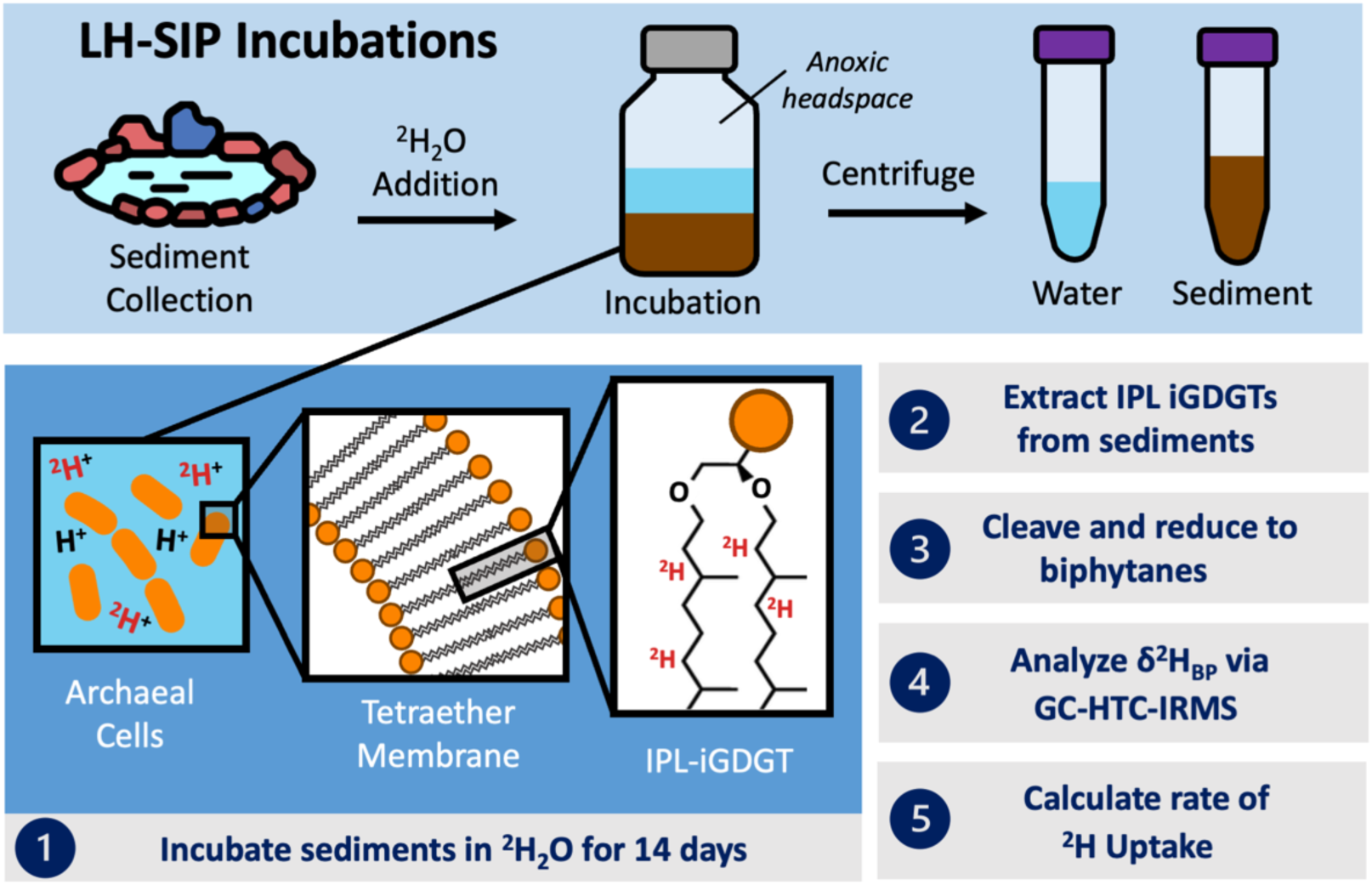
Schematic illustrating *ex situ* lipid hydrogen SIP assays performed on hot spring sediments. Whole sediments were incubated with 0.37 atom % ^2^H_2_O for 0, 3, or 14 days. Active archaeal cells are expected to incorporate the ^2^H label during *de novo* lipid synthesis (1). After incubations, IPL-iGDGTs are extracted from sediments (2), cleaved and reduced to component biphytane chains (3), and analyzed for their δ^2^H composition (4). The rate of ^2^H uptake into biphytane lipids is used to calculate apparent IPL-iGDGT-derived biphytane production rates and generation times (5).

The headspace of all serum bottles was then flushed with deoxygenated N_2_. Bottles were incubated at *in situ* temperatures (88 or 80 °C for Beryl and ‘ETAT-3’, respectively) for 0, 3, or 14 days in duplicate. Bottles were periodically shaken to evenly distribute the ^2^H_2_O labeling solution. At the end of the incubation, excess water was separated from sediments via centrifugation and collected for subsequent isotopic analysis. Sediment pellets were stored at -80 °C until lipid extraction.

To assess non-biological exchange or analytical artefacts and prevent false growth detections, two control treatments were also prepared. The live control contained sediments amended with unlabeled (e.g., natural abundance) filter-sterilized spring water. The killed control contained sediments that were sterilized via a freeze/autoclave cycle and amended with isotopically labeled spring water. Controls were incubated for 0 or 14 days with one replicate for each time point.

### 2.3 Water δ^2^H analysis

Aliquots of filter-sterilized natural abundance spring waters and labeled incubation waters (F_L_) were analyzed for δ^2^H at the Stable Isotope Laboratory at Dartmouth College via a dual inlet isotope ratio mass spectrometer (IRMS; Thermo Delta Plus XL) coupled to an H-Device for water reduction to H_2_ by chromium powder at 850 °C (Kopec et al., 2018). The δ^2^H_Water_ values were corrected using three laboratory standards spanning -161 to -7 ‰ vs. VSMOW, which were run every 8 to 10 samples. Labeled waters were first diluted (1:1000 w/w) with water of known isotopic composition to ensure they were in range of the internal standards. Analytical precision was <0.5 ‰.

Measured isotope values in δ^2^H (‰) notation on the VSMOW-SLAP scale were converted to F-values representing the absolute fractional abundance of ^2^H (unitless, but sometimes expressed as atom percent) using the isotopic composition of VSMOW [R_VSMOW_ = ^2^H/^1^H = 0.00015576] and the relationship: F = R/(1+R) = (δ + 1)/(1/R_VSMOW_ + δ + 1). After homogenization with pore water in the spring sediments, the isotopic composition of the labeled incubation water was diluted to 3,692 ± 218 ppm ^2^H (0.37 ± 0.02 atom %; ∼22,800‰) for Beryl, and 3,770 ± 278 ppm ^2^H (0.38 ± 0.03 atom %; ∼23,300‰) for ‘ETAT-3’. The diluted values, which represent a combination of tracer water and water present in spring sediments, were used for all growth rate calculations.

### 2.4 Lipid extractions and analyses

iGDGT lipids were extracted from each replicate using a Bligh-Dyer protocol for sediments (Lengger et al., 2012) with modifications for unique hot spring chemistry (Boyd et al., 2011). Total lipid extracts were treated with activated copper filings at room temperature overnight to remove S^0^ (Boyd et al., 2011). The intact polar lipid (IPL-iGDGTs) fraction was isolated from core lipids (core-iGDGTs) via elution in silica columns (Zhou et al., 2020). The IPL fraction was hydrolyzed in 5% HCl in MeOH at 70°C for 3 hours to remove polar headgroups and isolate the hydrocarbon core. Both IPL-derived-core-iGDGTs and free core-iGDGTs were identified and quantified via a UHPLC system coupled to a triple-quadrupole mass spectrometer (QQQ-MS) (Blewett et al., 2020; Zhou et al., 2020) at Harvard University, Cambridge, MA. The absolute abundance of iGDGT structures was determined by comparing the peak area of a co-injected C_46_-GTGT internal standard (Huguet et al., 2006).

A portion of the IPL-iGDGT fraction was further processed to release the component biphytane chains of iGDGT lipids, which is necessary for analysis of their δ^2^H composition via GC-HTC-IRMS. In brief, the ether bonds of iGDGTs were cleaved via digestion in hydroiodic acid and the resulting alkyl iodides were reduced to biphytanes (BPs) via hydrogenation in H_2_/PtO_2_. We refer to the resulting compounds as “iGDGT-derived BPs” throughout. Method evaluations indicate that the HI-H_2_/PtO_2_ procedure used here recovers approximately 67% of biphytane material, with similar relative recovery across BP-0, BP-1, and BP-2 (Rosendahl et al., 2026). Absolute recovery relative to the starting iGDGT pool cannot be determined because individual iGDGT structures yield different combinations of biphytane products during ether cleavage.

The identity and quantity of isoprenoid BP chains were analyzed via GC-MS (Thermo ISQ LT with TRACE 1310) for compound identification and via GC-flame ionization detector (GC-FID; Thermo TRACE 1310) for compound quantification at the University of Colorado at Boulder Earth Systems Stable Isotope Lab (CUBES-SIL, Boulder, CO). Biphytane absolute abundances were calculated using GC peak areas and co-injected n-alkane mixture as a quantification standard.

To describe the average number of cyclopentyl moieties (e.g., mean cyclization) in iGDGT and biphytane lipid distributions, we calculated Ring Indices for iGDGTs and biphytanes based on the relative abundances of specific compounds (Eq. 1, 2) (Zhang et al., 2016; Zhou et al., 2020). Crenarchaeol, an iGDGT that contains four cyclopentyl moieties and one cyclohexyl moiety, is treated as an iGDGT-4 in the RI_GDGT_ calculation. Crenarchaeol comprises < 0.4 % of the lipid pool for all samples and its contributions to RI_GDGT_ are minimal.

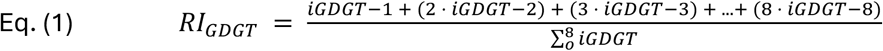

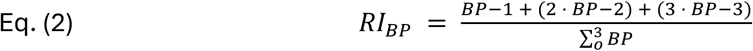

### 2.5 Biphytane δ^2^H analysis

The δ^2^H composition of individual BPs was determined by gas chromatography high temperature conversion isotope ratio mass spectrometry (GC-HTC-IRMS) on a GC IsoLink II IRMS System (Thermo Scientific) at the University of Colorado at Boulder Earth Systems Stable Isotope Lab (CUBES-SIL, Boulder, CO). The system consisted of a Trace 1310 GC fitted with a programmable temperature vaporization (PTV) injector and a 30 m DB5-HT column (250 μm inner diameter, 0.1 μm film thickness, Agilent, Santa Clara, CA, USA), ConFlo IV interface, and 253 Plus mass spectrometer (Thermo Scientific).

All ^2^H/^1^H ratios are reported in delta notation (δ^2^H, ‰) relative to VSMOW-SLAP.

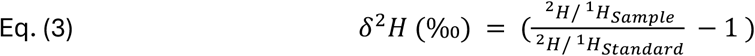

The apparent isotope fractionation between spring waters and BP lipids is reported in epsilon notation (^2^ε_L/W_, ‰).

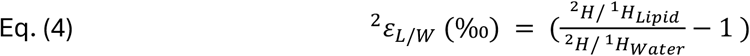

Hydrogen isotope values of individual biphytanes (δ^2^H_BP_) were measured relative to H_2_ reference gas and calibrated to the VSMOW scale using an A7 n-alkane standard (C_15_–C_30_, - 9 to -263 ‰ vs. VSMOW; A. Schimmelmann, Indiana University) and a C_36_ n-alkane (-259.2 ‰). Standards were run throughout each sequence at varying concentrations.

Data calibration was performed using 4022 compound-specific measurements obtained from 313 analyses of the combined standard mixture. The calibration workflow included corrections for background (bg) contribution, scale compression, and peak-size effects:

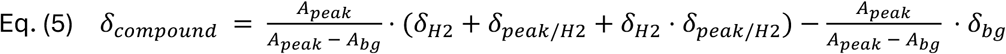

Peak areas represent time-integrated peak amplitudes after baseline subtraction, are H_3_ factor corrected, and range from 4 to 50 Vs. The reference gas (δ^2^H_2_) value was derived from the standard measurements and estimated based on the approach of Polissar and D’Andrea (2014). Two different hydrogen tanks were used during the analyses: (i) tank 1 (2024) had a δ^2^H_2_ of -312 ± 2 ‰ and (ii) tank 2 (2025) had a δ^2^H_2_ of -259 ± 3 ‰. The uncertainty estimates for individual analytes were defined conservatively by selecting the largest of the following three metrics: (i) the propagated calibration uncertainty (0.3 to 1.8 ‰), (ii) the bias-corrected pooled standard deviation of the samples (2.1 to 15.8 ‰) or (iii) the bias-corrected standard deviation of the nC36 standard (3.5 ‰). The bias-correction and standard deviation pooling was conducted following the approach of Polissar and D’Andrea (2014). Additionally, analyte δ^2^H values were corrected for the addition of H during the alkyl iodide hydrogenation by Pt/H2 following the procedure reported by Rosendahl et al. (2026). The resulting hydrogenation correction systematically shifted the δ^2^H values of the alkyl chains by +5.5 to +14.5 ‰ and increased analytical uncertainty by up to 2.5 ‰. For each sample, corrected δ^2^H_BP_ values were averaged across analytical replicates (n ≥ 3) and biological duplicates, where available, using weighted means (1/σ^2^). Reported errors represent propagated uncertainty. Abundance-weighted means include all BPs with > 5 % relative abundance.

### 2.6 Growth rate, apparent generation time, and lipid biosynthesis rate calculations

Biphytanes are hydrocarbon chains that incorporate H during biological activity and contain only C-H bonds that are nonexchangeable over biological time scales (Schimmelmann et al., 2006). The incorporation of the ^2^H label into biphytanes is calculated as follows (Kopf et al., 2016; Caro et al., 2023):

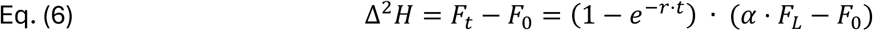

Where *r* is the specific biosynthesis rate (1/day), *t* is the duration of the incubation (days); α is the assimilation efficiency and fractionation of water hydrogen during lipid biosynthesis; and *F_0_*, *F_t_*, and *F_L_* are the fractional abundances of ^2^H in biphytanes at time 0, biphytanes at time *t*, and in the isotopically labeled spring water, respectively. Assimilation efficiency, α, represents the fraction of iGDGT-bound hydrogen sourced from water (as opposed to the organic substrate used for growth, which also contributes H to lipids during heterotrophic growth).

For a given community of archaea, assimilation efficiency will depend on the dominant metabolism. Lab experiments on a thermoacidophilic archaeon, *Sulfolobus acidocaldarius,* grown heterotrophically on simple sugars revealed assimilation values of 0.56 ± 0.01 (Harris et al., 2025). Similar experiments on a thermophilic archaeon, *Archaeoglobus fulgidus*, grown autotrophically using CO_2_ as the carbon source, H_2_ as the electron donor, and thiosulfate as the terminal electron acceptor, revealed assimilation values of 0.76 (Rhim et al., 2024). This study also cultured *A. fulgidus* heterotrophically and determined α > 0.50. Here, we consider the range of potential assimilation efficiencies for Archaea growing with different metabolisms by setting the mean α-value to 0.66 and its error (σ_α_) to 0.1, which encompasses a purely heterotrophic community (α = 0.56) and a purely autotrophic community (α = 0.76).

Because lipid biosynthesis represents both growth and repair processes, the calculated *r* value (days^-1^) provides an upper bound for the specific growth rate, *µ*, and a lower bound for the apparent generation time, *T_G_* (days), of the Archaea that produce a certain BP.

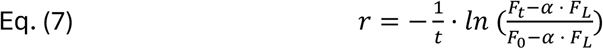

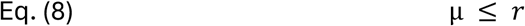

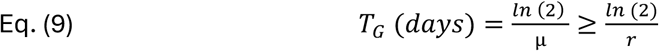

Based on the apparent generation time and a representative estimate of the initial standing stock of IPL-derived biphytanes (B_0_) in sediments from each spring, we estimate the maximum annual lipid production rate (P), in units of ng BP g^-1^ dry sediment year^-1^, as follows:

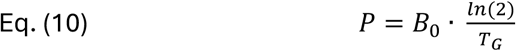

where *T_G_* is expressed in years. These equations provide BP-specific growth rate, apparent generation time, and annual production rate estimates which can be aggregated into an abundance-weighted mean using the relative abundance of each BP moiety. The aggregated mean represents an abundance-weighted estimate of apparent lipid biosynthesis for the pooled iGDGT-producing archaeal community. Under steady-state conditions, where lipid production and degradation are approximately balanced and the standing stock remains constant through time, the apparent generation time is equivalent to the turnover time of the sedimentary lipid pool (see Supplementary Information S3 for detailed calculations).

Considering the inputs and their errors for our *ex situ* LH-SIP incubations, a minimum ^2^H uptake (i.e., Δ^2^H) of 1.5 ppm (∼9.6 ‰, VSMOW) is necessary to detect biosynthesis at the 2σ level (Figure S5). Over a 14-day incubation, this corresponds to a minimum detectable growth rate of 0.017 ± 0.008 year^-1^ and a maximum detectable apparent generation time of 42 ± 21 years. Given the constraints of our incubations (e.g., F_L_ = ∼ 0.37 atom % ^2^H; t = 3, 14 days), we performed a sensitivity analysis across a realistic range of Δ^2^H enrichments from 1.56 ppm to 1556 ppm (corresponding to +10 to +10,000 ‰, respectively), and demonstrate that our SIP incubations can detect archaeal generation times ranging from 2 days to 42 years (Figure S6). See Supplementary Information S1, S2 for detailed calculations.

All statistical analyses were performed in R version 4.3.1 using RStudio (RStudio Team, 2023). Raw data and code for all data manipulations and statistical analyses is available on Zenodo (Harris, 2026).

## 3 Results

### 3.1 Spring geochemistry and archaeal lipid inventory

Beryl and ‘ETAT-3’ are circumneutral hot springs characterized by high temperatures (approaching the boiling point in each system), suboxic (<0.7 mg L^-1^ O_2_), and reducing redox conditions (Beryl: -334 mV, ‘ETAT-3’: -84 mV) (Table 1, Figure 1). This suggests that oxygen penetration into sediments is minimal at the sampled locations. ‘ETAT-3’ has substantially higher specific conductance than Beryl (23.5 vs. 2.5 mS cm^-1^) reflecting the high evaporation potential of the hyperarid Atacama Desert and/or contact with brines (Risacher et al., 2003), and the dilution of Beryl hydrothermal water with meteoric water (Truesdell, 1977). The δ^2^H composition of spring waters differs between the sites (Beryl: -139 ‰; ‘ETAT-3’: -68 ‰), reflecting global hydrology patterns and regional evaporation trends (Hurwitz and Lowenstern, 2014; Munoz-Saez et al., 2018; Harris et al., 2025; Truesdell, 1977). Both sites exhibited large negative apparent ^2^H fractionation between BP lipids and spring waters, although fractionation was more negative at Beryl (−309 ‰) than at ‘ETAT-3’ (−255 ‰) (Table 1).

At each site, both IPL- and core-iGDGTs were recovered from sediments with pooled concentrations of 80 and 12 µg iGDGT per g dry sediment for Beryl and ‘ETAT-3’, respectively (Table 1, Figure 3). The IPL contribution to the total iGDGT pool was 15 and 88 % for Beryl and ‘ETAT-3’, respectively. iGDGTs with 0 to 7 cyclopentane moieties (i.e., iGDGT-0 to iGDGT-7) were detected in sediments at both sites, with the less ringed moieties (iGDGT <5) dominating both IPL and CL distributions (Figure 3, Table S1). iGDGT ≥ 5 represented < 2 % of the IPL-iGDGT pool and 6-8 % of the core-iGDGT pool at both sites. In Beryl sediments, iGDGT-0 to -4 each contributed ∼20 % to the IPL lipid pool, corresponding to a RI_GDGT_ of 2.06 ± 0.01. When cleaved and reduced to biphytane lipids (BPs), BPs -0, -1, and -2 each contributed ∼30 % to the Beryl IPL lipid pool, corresponding to a RI_BP_ of 0.97 ± 0.01. ‘ETAT-3’ sediments were dominated by iGDGT-2 to iGDGT-4 (20, 20, and 30 %, respectively), corresponding to a RI_GDGT_ of 2.73 ± 0.05 (Figure 3). BP-0, -1, and -2 contributed ∼20, 30, 40% to the IPL pool, corresponding to a RI_BP_ of 1.08 ± 0.12. At both sites, BPs ≥ 3 represented < 3% of the BP pool.

**Figure 3.**
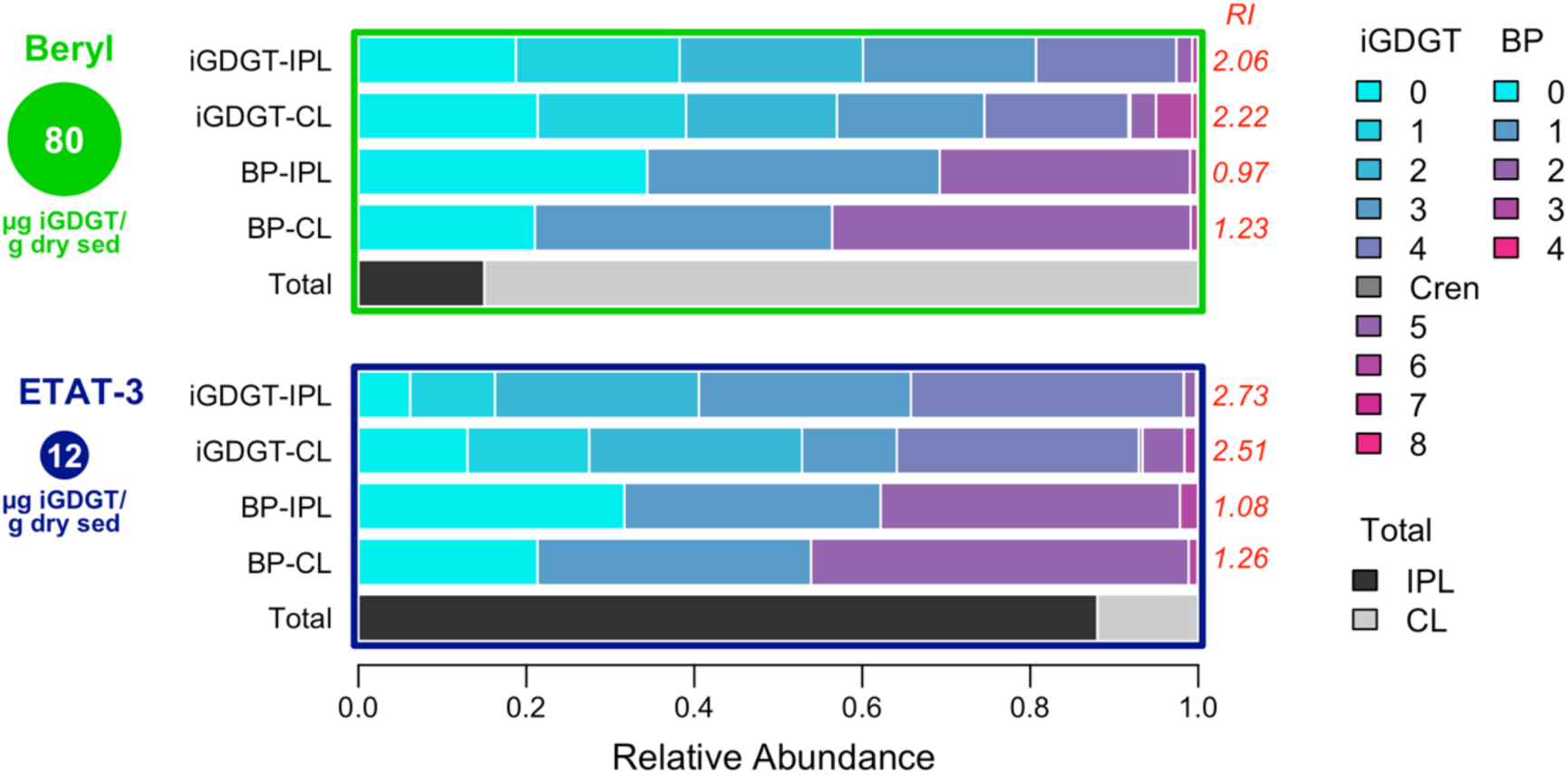
Archaeal lipid inventories and compositions. Bars represent the relative composition of archaeal intact polar iGDGTs (iGDGT-IPL) and core iGDGTs (iGDGT-CL) and corresponding iGDGT-derived biphytane (BP) lipids. The red numbers to the right of each bar indicate Ring Indices (RI) calculated for iGDGT or BP lipids. The total bar depicts the relative proportion of IPL vs. core lipids in the cumulative sedimentary iGDGT pool. The circles underneath the site names are scaled to represent the total sedimentary iGDGT inventory in µg iGDGT per g dry sediment.

### 3.2 LH-SIP experimental assays

The relative abundance of specific iGDGT and BP moieties in the IPL pool for experimental and control replicates was monitored over the 14-day SIP experimental assays. No changes in lipid distributions or related Ring Indices were observed during the incubation (Figure S2, Table S1). In unlabeled control samples, the δ^2^H composition of IPL-iGDGT-derived biphytanes was -405 ± 2 ‰ at Beryl and -306 ± 1 ‰ at ‘ETAT-3’ (Figure S3, Table S1). These values are within the natural variation observed for biphytanes in hot spring sediments in Yellowstone and El Tatio (Harris et al., 2025). The δ^2^H composition of biphytanes in the killed controls also remained within this natural-abundance range, indicating no detectable non-biological incorporation of the isotope label during the incubation (Figure 4, Table S1).

**Figure 4.**
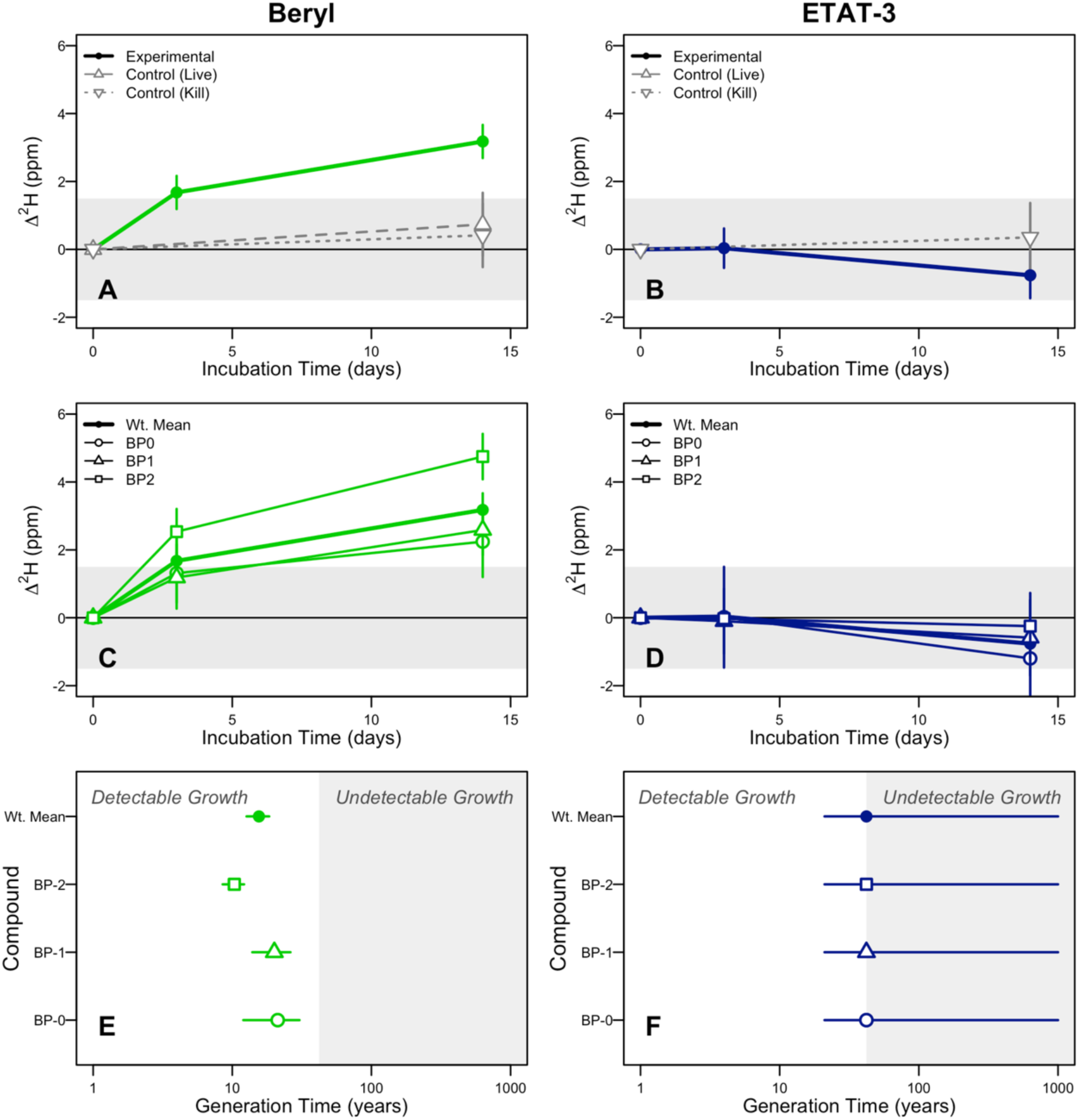
The Δ^2^H enrichment, defined as the change in ^2^F (ppm) relative to the start of the incubation, of archaeal IPL-iGDGT-derived biphytane lipids over the course of LH-SIP incubations for (A, B) experimental and control samples averaged across biphytane moieties, and for (C, D) individual biphytane moieties. ^2^H-uptake into biphytanes was observed after 3 and 14 days in Beryl sediments; no uptake was observed in ‘ETAT-3’ sediments. Associated compound-specific and aggregated (abundance-weighted mean) apparent generation times (E) were calculated for Beryl using the integrated Δ^2^H between 0 and 14 days. Apparent generation times indicated for ‘ETAT-3’ (F) represent the minimum generation time (∼42 years) consistent with the lack of ^2^H-uptake observed (see detection limits and sensitivity analysis in Figures S5, S6). The shaded regions indicate ^2^H-uptake (A-D) or apparent generation times (E,F) that are unresolvable by our LH-SIP incubations. Points represent means across biological replicates; error bars indicate propagated error from biological replication, technical replication, and uncertainty in assimilation efficiencies (for generation time calculations). Note: if archaea recycle or re-functionalize pre-existing lipid components during membrane synthesis, these calculated apparent generation times may overestimate true cellular generation times.

To determine apparent lipid biosynthesis rates and generation times, the uptake of the ^2^H_2_O labeling solution into lipids over the 14-day incubation was determined. At Beryl, we observed minor uptake of the ^2^H-label into biphytane chains after 3 days (Δ^2^H = 1.7 ± 0.5 ppm; 11 ± 3 ‰) and a larger enrichment after 14 days (Δ^2^H = 3.2 ± 0.5 ppm; 20 ± 3 ‰) (Figure 4A, Table 2). When integrated across the 3- and 14-day measurements, the abundance-weighted average apparent generation time was 16 ± 3 years, corresponding to an annual lipid production rate of 32.0 ± 5.4 ng BP g^-1^ dry sediment year^-1^ for the actively cycling IPL-derived biphytane pool. Apparent generation time varied slightly with BP ring number, with more cyclized biphytanes showing shorter generation times. BP-0, -1, and -2 have generation times of 21, 20, and 10 years, respectively, corresponding to annualized lipid production rates of 25.5, 27.6, and 44.7 ng BP g^-1^ dry sediment year^-1^ (Figures 4C, E, S4; Table 3).

**Table 2.**
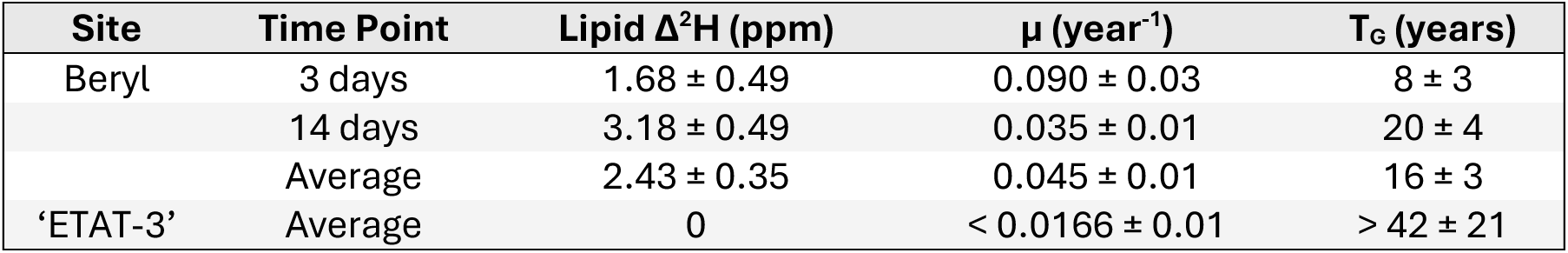
The aggregated Δ^2^H enrichment (ppm) for iGDGT-derived biphytane lipids relative to the start of the incubation, averaged across all biphytane moieties. The enrichment was used to calculate aggregated growth rates (μ, year^-1^) and apparent generation times (T_G_, years) for sedimentary archaeal lipids. For Beryl, values are shown for individual time points and integrated over 3- and 14-day time points. For ‘ETAT-3’, which showed no ^2^H incorporation into lipids over 14 days, the detection limits of our SIP incubations are shown; these limits represent the maximum growth rate and minimum apparent generation time consistent with the lack of uptake. Error in Δ^2^H is propagated across biological and technical replicates. Error in growth rate and apparent generation time also accounts for the propagated uncertainty of isotopic measurements and tracer assimilation efficiency and variation across biological replicates.

**Table 3.** Compound-specific growth rates (μ, year^-1^), apparent generation times (T_G_, years), and lipid production rates (ng biphytane g^-1^ dry sediment year^-1^) for Beryl sedimentary archaeal lipids, integrated over 3- and 14-day time points. Also shown are calculations of estimated new biomass production (ng biphytane g^-1^ dry sediment) after 14-days and the increase in IPL biphytane biomass relative to the start of the incubation. Error accounts for variation among biological and technical replication and the propagated uncertainty of isotopic measurements and tracer assimilation efficiency. (See Supplementary Information S3 for an explanation of calculations.)

| Compound | $\mu$ (year<sup>-1</sup>) | $T_G$<br>(years) | Production<br>rate (ng g<sup>-1</sup><br>sed year<sup>-1</sup>) | New<br>biomass<br>production<br>(ng g<sup>-1</sup> sed) | % Biomass<br>increase |
| --- | --- | --- | --- | --- | --- |
| BP-0 | 0.033 ± 0.01 | 21 ± 9 | 25.5 ± 11.1 | 0.98 ± 0.43 | 0.13 ± 0.05 |
| BP-1 | 0.035 ± 0.01 | 20 ± 6 | 27.6 ± 8.4 | 1.06 ± 0.32 | 0.13 ± 0.04 |
| BP-2 | 0.067 ± 0.01 | 10 ± 2 | 44.7 ± 7.9 | 1.72 ± 0.30 | 0.26 ± 0.05 |
| Wt. Average | 0.044 ± 0.01 | 16 ± 3 | 32.0 ± 5.4 | 1.23 ± 0.21 | 0.17 ± 0.03 |

At ‘ETAT-3’, no ^2^H incorporation was detected above the assay detection limit, constraining the abundance-weighted apparent mean generation time to > 42 ± 21 years and the maximum lipid production rate to ≤0.017 ± 0.008 µg BP g^-1^ dry sediment year^-1^ for the actively cycling IPL-derived biphytane pool (Figures 4B, D, F, S4; Table 2). We observed no uptake of the ^2^H-label into biphytanes in control treatments at either hot spring.

Although an increase in absolute IPL-BP abundance over time may provide evidence for *de novo* production, the standing IPL pool at each spring is sufficiently large that the estimated new production during the 14-day incubation (<0.3% of standing pool; Table 3) would be unresolvable via our GC-MS methods.

## 4 Discussion

### 4.1 Lipid biosignatures indicative of live sedimentary Archaea

While iGDGTs are virtually ubiquitous in marine and terrestrial systems, the most highly ringed moieties (e.g., iGDGT-7 and -8) have only been detected in hydrothermal systems (Schouten et al., 2007). Here, we detect iGDGTs-0 to -7 in the core and IPL fraction of both hot springs; the presence of higher ringed moieties suggests autochthonous production, as opposed to allochthonous inputs from surrounding soils (Boyd et al., 2013). Although IPLs, which contain intact, functional head groups, are often interpreted as indicators of living or recently synthesized biomass, increasing evidence suggests that some IPLs may persist in sediments for years to millennia under low-energy conditions. Core-iGDGTs are known to accumulate and persist long after cell death and are interpreted as fossil lipids (Schouten et al., 2013; Chong, 2024). The presence of IPL-iGDGTs in the sediments of both hot springs suggests that live archaea were present. Although Beryl contained a larger total iGDGT pool, IPL-iGDGTs comprised a substantially greater fraction of the total iGDGT pool at ‘ETAT-3’ than at Beryl (88% vs. 15%). As a result, the absolute inventories of IPL-iGDGTs were similar between the two sites (12 and 11 µg g^-1^ dry sediment at Beryl and ‘ETAT-3’, respectively).

### 4.2 Activity of sedimentary Archaea in SIP incubations

We monitored several parameters during SIP incubations that can indicate archaeal activity, including ^2^H-label uptake into iGDGT-derived biphytanes and changes in the relative abundance of specific lipid compounds. After 14-day incubations, the abundance-weighted mean δ^2^H composition of BP lipids at Beryl increased by ∼ 18 ‰, corresponding to a lower-bound apparent generation time of 16 ± 3 years aggregated across biphytane moieties (Figures 5, S2). More highly cyclized biphytanes also showed systematically shorter apparent generation times, with BP-2 exhibiting faster apparent turnover than BP-0 and BP-1 (Figure S4). This pattern may reflect preferential synthesis of highly cyclized membrane lipids by metabolically active Archaea under hydrothermal conditions. No ^2^H-uptake was observed in ‘ETAT-3’ sediments, indicating new lipid biosynthesis was below the sensitivity of the assay, which could resolve apparent generation times up to 42 ± 21 years. To our knowledge, these data provide the first LH-SIP constraints on archaeal IPL-iGDGT production rates in terrestrial hot spring sediments.

**Figure 5.**
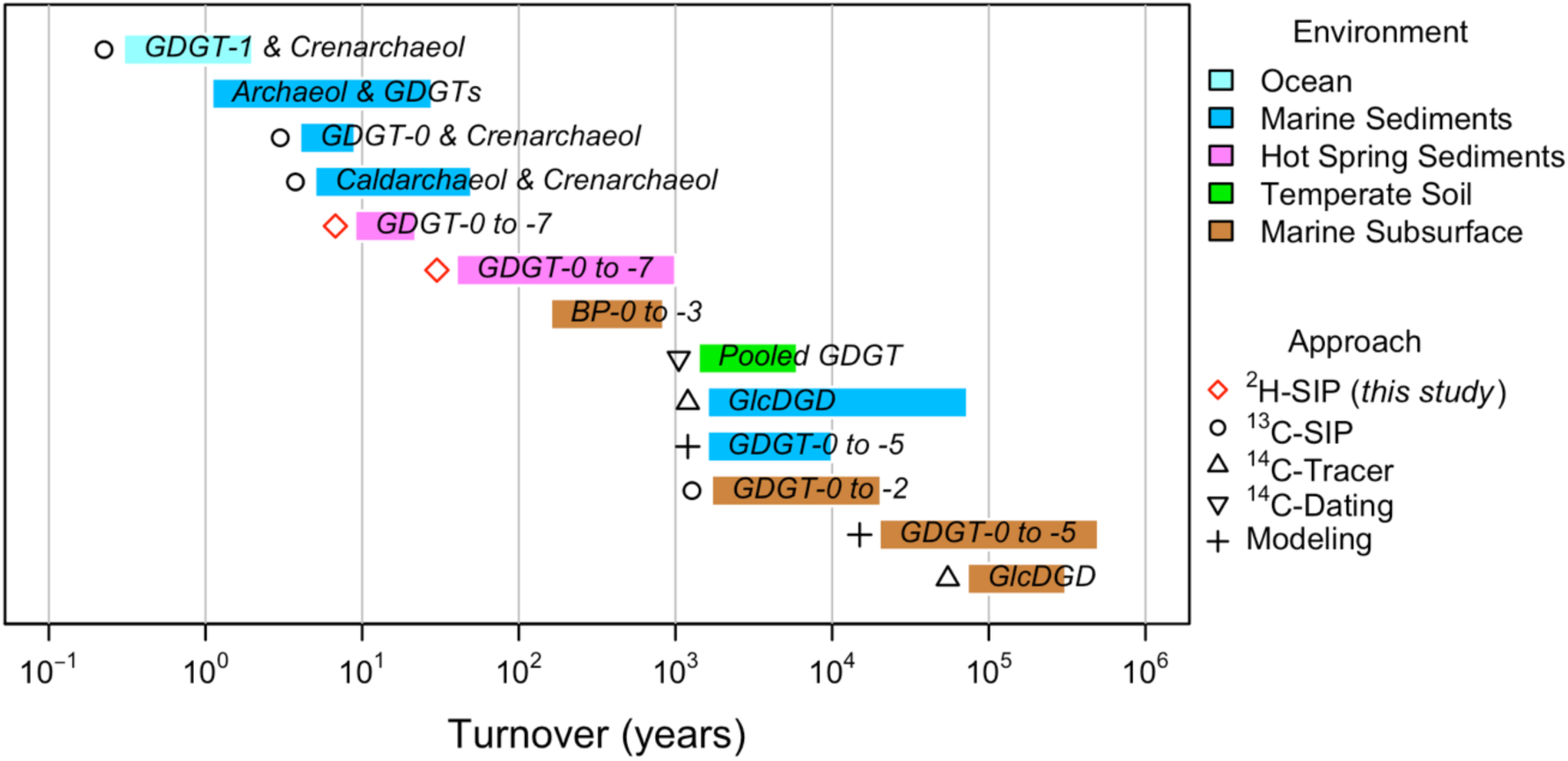
Reported turnover and generation times for a variety of archaeal lipids in marine and terrestrial environments including ocean waters, marine surface sediments, terrestrial soils, and the deep marine subsurface (indicated by bar color). Red diamonds indicate the results of this study. Reported turnover time ranges from weeks (ocean waters) to hundreds of thousands of years (deep marine biosphere). These estimates were reported in a variety of terminology and units, including growth rate, generation time, turnover rate, turnover time, and doubling time. For consistency, all published estimates were converted to turnover time (years). In this framework, turnover time and generation time are mathematically equivalent, but turnover time explicitly assumes steady-state biomass conditions in which production and loss rates are balanced. All lipids included are archaeal in origin, primarily tetraethers GDGT-0 through GDGT-5 and crenarchaeol. GlcDGD is an archaeal diether, comprising a glucosyl (sugar) headgroup attached to a diphytanylglycerol diether core. Methods (indicated by symbol shape) include stable isotope probing (SIP) with ^13^C or ^2^H tracers, ^14^C radiotracer or dating methods, and modeling approaches. This compilation highlights the wide variability in archaeal lipid turnover, which spans six orders of magnitude across environments. Full metadata for each entry are provided in Table S2.

These apparent rates are substantially slower than expected based on the limited previous *in situ* estimates of microbial generation times for planktonic cells in the water column of circumneutral/alkaline springs (Bott and Brock 1969; Brock et al., 1971; Bohlool and Brock, 1974; Mosser et al., 1974), and substantial microbial activities documented in sediments of such springs (Urschel et al., 2015; Fernandes-Martins et al., 2023). These observations suggest that the decadal-scale apparent generation times measured here may overestimate true cellular generation times. The discrepancy between the decadal-scale apparent generation times measured here and independent observations of microbial activity suggests that additional processes may decouple BP turnover from cellular growth.

One important alternate explanation is that a substantial fraction of the sedimentary IPL-iGDGT pool may persist for much longer than the actively growing cells that originally produced it. Although IPL-iGDGTs are often interpreted as markers of living or recently synthesized biomass, preserved IPLs from inactive or dead cells would not incorporate the ^2^H_2_O label. A large standing pool of such material would dilute the isotopic enrichment of newly synthesized lipids and make the calculated apparent generation times substantially longer than the true generation times of active archaeal cells. Notably, IPL-iGDGTs comprised a much larger fraction of the total iGDGT pool at ‘ETAT-3’ than at Beryl (88% vs. 15%). If a substantial portion of this IPL pool represents persistent relict material, this may contribute to the lack of observed ^2^H incorporation at ‘ETAT-3’.

A related possibility is that some fraction of the sedimentary IPL pool is allochthonous rather than produced by benthic sedimentary archaea *in situ*. However, the highly cyclized iGDGT distributions observed here, together with previous evidence that iGDGT composition and abundance in Yellowstone hot springs are distinct from surrounding soils (Boyd et al., 2013), strongly support *in situ* hot spring production rather than substantial input from non-hydrothermal terrestrial sources. Still, some of these lipids could have been produced by archaeal cells in the spring water column and subsequently deposited to the sediments. Hot spring waters can host diverse free-living archaeal plankton, even in visually clear and low-biomass systems (Reysenbach and Cady, 2001; Colman et al., 2016; Fernandes-Martins, 2021). Although sedimentation is likely less efficient than in marine systems, deposition of suspended cells or their remnants is possible, particularly in springs with deeper basins, reduced flow, or longer water residence times. Even in higher-flow systems, low-energy boundary layers at the sediment–water interface could facilitate accumulation of suspended biomass. If water-column-derived IPL-iGDGTs accumulate in the sediments but are no longer actively synthesized after deposition, they would enlarge the standing background lipid pool and dilute ^2^H incorporation by benthic cells. Because particulate lipids in the overlying water column were not quantified here, the potential contribution of water-column or suspended archaeal biomass remains unresolved.

More broadly, it is likely that some archaeal lipids in these sediments are produced through partial recycling or salvage of existing lipid components, rather than by wholly *de novo* synthesis. For example, biphytanyl chains could be recycled from relict archaeal membranes or detrital core lipids, in which case only the glycerol backbone or polar headgroups of iGDGTs are newly assembled (Takano et al., 2010; Fehyl-Buska, 2016). Archaeal culture studies have demonstrated the potential for isoprenoid salvage pathways in some thermophilic archaea (e.g., *Sulfolobus acidocaldarius*; Ohnuma et al., 1996). While this process could theoretically proceed via hydrolysis and reformation of the ether bond to the glycerol backbone, ether cleavage imparts a steep energy cost (∼360 kJ mol^−1^) (White et al., 1996). If recycling occurs, active membrane synthesis could occur with minimal ^2^H incorporation into biphytane chains, causing our BP-specific measurements to underestimate lipid production rates, and overestimate apparent generation times.

Such recycling was proposed for archaea living in deep sea sediments, where *ex situ* SIP incubations showed that ^13^C-substrates were incorporated into glycerol units at a faster rate than isoprene units, indicating faster head group synthesis than biphytane synthesis (Takano et al., 2010; Liu et al., 2011). Culture studies have since confirmed biphytane recycling within certain iGDGTs (Lin, 2009) and this process has been observed in surface sediments of a hypoxic marine lake (Lipsewers et al., 2018), evaporitic facies of Dead Sea deep sediments (Thomas et al., 2019), surface and subsurface deep sea sediments (Lipp and Hinrichs, 2009), and the Cathedral Hill deep sea hydrothermal vent in Guaymas Basin (Bentley et al., 2022). Based on these observations, it is hypothesized that lipid recycling may be an adaptive mechanism for archaea living in low-energy environments to minimize energy expenditure on growth and maintenance. Hot springs can also be energy-limiting environments, particularly those with acidic, anoxic, or low organic carbon conditions (Des Marais and Walter, 2019), suggesting recycling may occur in hydrothermal systems as well. Additional research, perhaps following the ^13^C-sugar SIP methods of Lin and others (2013), is needed to quantify the extent to which archaea in terrestrial hot springs recycle exogenous core lipids into their own cellular membranes.

Similarly, ^2^H incorporation into biphytanes may lag cellular growth if iGDGT biosynthesis involves precursor pools, delayed head-to-head condensation of diether intermediates, or post-synthetic remodeling of head groups. For example, in archaeal IPL-SIP experiments with anaerobic methanotrophs (ANME-1) enrichments, ^2^H-label uptake into tetraethers lagged substantially behind uptake into diether lipids, likely causing short incubations to underestimate tetraether production rates (Kellermann et al., 2016). Although the biosynthetic pathways of thermophilic hot spring archaea may differ, similar delays would bias our 14-day incubations toward underestimating true iGDGT production rates.

Finally, bottle effects have been observed in incubations in acidic and alkaline springs in Yellowstone (Boyd et al., 2009; Keller et al., 2023), so it is also possible that the long apparent generation times we observed are due to the experimental conditions. To maximize environmental fidelity, incubations were performed on whole sediments at *in situ* temperatures under anoxic conditions using filtered natural spring water as the labeling solution. No additional electron donor/acceptor or carbon sources were provided. Sediments experienced several days of transport after field collection, so transport and handling could have altered microbial activity to some degree. Accordingly, we consider it unlikely that incubation artifacts alone can explain the extremely long apparent generation times observed here.

Overall, our LH-SIP results indicate that turnover of sedimentary IPL-iGDGT-derived biphytanes is substantially slower than expected in these hot spring communities. The apparent decadal-scale rates likely reflect substantial dilution by pre-existing IPLs and recycling or re-functionalization of existing lipid components, in addition to *de novo* lipid synthesis. Thus, the LH-SIP measurements are best interpreted as constraints on sedimentary biphytane turnover rather than direct estimates of archaeal cell generation times.

### 4.3 Comparison to archaeal iGDGT production rates in other environments

Measurements of iGDGT production in hydrothermal environments are rare, but several SIP experiments have been performed to determine the production rate of IPL-iGDGT lipids in other environments (Figure 5, Table S2). To facilitate comparison across studies using different experimental approaches and terminology, all reported rates were converted to equivalent turnover times (years), with turnover implicitly assuming steady-state biomass conditions. SIP experiments targeting Thaumarchaeota in the North Sea water column report relatively rapid turnover, ranging from 70 to 130 days for crenarchaeol (Wuchter et al., 2003; Pitcher et al., 2011). Experiments with ^14^C-bicarbonate and ^14^C-acetate in shallow sediments of the Mediterranean Sea revealed iGDGT turnover times of ∼3 to 24 years depending on experimental conditions (Evans et al., 2019). In marine sediments in Iceland, ^13^C-substrate SIP experiments suggest similar turnover, with estimated crenarchaeol and iGDGT-0 generation times of at least 4 to 9 years (Lengger et al., 2014). Much longer turnover times ranging from 1,700 to 20,500 years were observed for iGDGTs-0 to -2 in anoxic marine surface sediments during 468-day SIP incubations with various ^13^C-labeled compounds (Lin et al., 2013). This extremely slow uptake, even when amended with labile carbon sources, suggests minor turnover of sedimentary iGDGTs (Lin et al., 2013).

Degradation experiments using ^14^C-labeled diether lipids in low energy conditions in the marine subsurface report estimated half-lives of up to 310 kyr (Xie et al., 2013). These studies suggest that in deep subsurface environments, IPL-iGDGTs may persist for thousands to hundreds of thousands of years, indicating that a significant fraction represents fossil material rather than active biomass.

In comparison, our LH-SIP experiments indicate apparent IPL-iGDGT-derived BP turnover on decadal timescales at Beryl (16 ± 3 years), while turnover at ‘ETAT-3’ was too slow to resolve under the conditions of this assay (> 42 ± 21 years; inferred from assay detection limits). These apparent turnover times are similar in magnitude to reports from surface marine sediments (Lengger et al., 2014; Evans et al., 2019) but should not be interpreted as direct cellular generation times based on the likelihood of a persistent standing pool of IPL-iGDGTs and the potential for lipid recycling or re-functionalization in hot spring sedimentary archaeal communities.

### 4.4 Considerations for future LH-SIP applications to hot spring sediment archaeal communities

Our detection of ^2^H uptake into biphytanyl chains of hot spring sedimentary Archaea in 14-day incubations highlights the utility of the ^2^H_2_O SIP approach to polyextreme environments, characterized by archaeal inhabitants with low rates of lipid turnover. In future studies targeting environments with similarly extreme conditions, this approach could better resolve slow biosynthesis rates by using a more enriched labeling solution (e.g., 5 to 10 atom % ^2^H) (Figure S5), though labeling solutions above 20 atom % are not recommended because of the potential for ^2^H_2_O toxicity (Berry et al., 2015; Kopf et al., 2016). Longer incubations (e.g., months or longer) will also permit detection of slower lipid production, though there is a tradeoff between short incubations that more naturally reflect *in situ* conditions and longer incubations, which increase sensitivity to slow ^2^H uptake but may be less representative of natural conditions.

Future LH-SIP studies could also improve environmental fidelity by initiating incubations (e.g., adding ^2^H-tracer) immediately after sediment collection, thereby minimizing the time interval between sampling and isotope amendment. In field settings with appropriate permitting and logistics, sealed incubation vessels or continuous flow vessels (Boyd et al., 2009) could also be deployed directly within a hot spring, allowing SIP experiments to proceed under more natural thermal conditions and reducing potential artifacts associated with sample transport.

Additionally, the ^2^H_2_O-SIP assay used here could be coupled with ^13^C-labeled substrates, which has several potential benefits. Amending *ex situ* incubations with ^13^C-dissolved inorganic carbon permits the distinction between autotrophic and heterotrophic growth (Wegener et al., 2012). Also, ^13^C-organic substrates can potentially stimulate growth by slow-growing microbes, which can increase the likelihood of detecting microbial activity, though the resulting lipid synthesis rates should not be considered representative of *in situ* dynamics (Wegener et al., 2016).

Future studies investigating iGDGT production rates in hot spring sediments would also benefit from knowledge of the proportion of IPL-iGDGT pools that represent living vs. fossil biomass. If hot spring conditions favor preservation of IPL-iGDGTs, as do conditions in the deep biosphere, then this large background signal could obscure isotope label incorporation into newly synthesized lipids, resulting in erroneously slow biosynthesis rates. This context is particularly important to consider when investigating whole sediments, where suboxic to anoxic conditions and protective sediment mineral matrices likely slow degradation rates (Reysenbach and Cady, 2001; Kaur et al., 2011; Parenteau et al., 2014).

Age dating (e.g., radiocarbon dating) provides one approach to quantify the active vs. fossil IPL pool. IPL lipids with similar ages to core lipids would suggest preservation of IPL lipids, whereas younger IPLs would suggest recent synthesis. Another way to isolate a younger fraction of the IPL pool is to separate IPL-iGDGTs based on the structure of their polar head groups – in marine environments lipids with glyco-head groups have been shown to be preserved over much longer timescales than lipids with phospho-head groups (Lipp and Hinrichs, 2009; Logemann et al., 2011; Xie et al., 2013). High-resolution LC-MS characterization of intact IPLs could further resolve the diversity and relative abundance of specific phospho-, glyco-, and mixed headgroup structures present before hydrolysis, providing additional context for interpreting apparent lipid turnover. Pairing this structural information with LH-SIP measurements of individual IPL classes could help determine whether low ^2^H-incorporation reflects genuinely slow synthesis or dilution by more persistent headgroup-specific lipid pools. Examining phospho- and glyco-headgroup-bearing iGDGTs separately would therefore help constrain the relative sizes of active and fossil iGDGT pools.

Future studies could target other lipid classes in addition to iGDGTs, for example archaeal diethers or bacterial fatty acids. Measuring ^2^H incorporation into archaeol-derived phytane alongside iGDGT-derived biphytanes would provide a useful comparison of diether and tetraether lipid synthesis. Because archaeol and related diether lipids may serve as precursors or intermediates in tetraether lipid biosynthesis, differences in their labeling rates could help distinguish genuinely slow archaeal growth from delayed iGDGT synthesis, remodeling, or recycling of pre-existing lipid components. Previous SIP experiments have shown that isotope incorporation into archaeal tetraether lipids can lag behind incorporation into diethers, suggesting that paired measurements of these lipid classes could help identify biosynthetic delays that bias short-term iGDGT-based growth estimates (Kellermann et al., 2016).

Finally, the LH-SIP assay used here returns biosynthesis rates that are aggregated across all cells that produce a certain BP lipid structure. Variation in growth rate among different archaeal taxa, taxa with different metabolisms, or among individual cells in natural systems is unresolved, although genome replication estimation techniques may hold promise in eventually overcoming these limitations (e.g., Fernandes-Martins, 2021). In addition, variation among specific iGDGTs is obscured because the eight iGDGT structures detected are cleaved and reduced to four component biphytane structures (though this could be solved with preparatory separation via liquid chromatography). This lack of specificity presents a challenge, particularly in systems where cells may be distributed patchily and/or growth rates may be heterogeneous among metabolic strategies and environmental micro-niches. As such, employing single-cell methods, such as SIP-NanoSIMS, which can resolve cell-specific growth rates and µm-scale variation in background isotopic features (Berry et al., 2015; Eichorst et al., 2015; Trembath-Reichert et al., 2017; Caro et al., 2024) may more accurately resolve archaeal growth rates in hot spring sediments.

### 4.5 Relevance to Mars Analogs and Mars Exploration

Our work provides empirical constraints on archaeal lipid production under conditions relevant to hydrothermal deposits on Mars and supports the inclusion of lipid-focused biosignature detection strategies in future Mars missions and sample return campaigns. Hydrothermal environments are among the most promising astrobiological targets on Mars because these systems were potentially capable of sustaining microbial life when active and have the potential to preserve biosignatures over geologic timescales (Cady et al., 2018; Finkel et al., 2023). Mineral deposits associated with ancient hydrothermal activity, including silica sinters and phyllosilicate-rich outcrops, have been identified at sites such as Nili Patera, Arabia Terra, Gusev crater, Valles Marineris, and Jezero crater (Rossi et al., 2008; Skok et al., 2010; Ruff et al., 2020; Rogers et al., 2023; Beck et al., 2025). The terrestrial hot springs studied here provide valuable analog systems for understanding microbial habitability and biosignature preservation in these Martian environments (Cady et al., 2018; Teece et al., 2020).

The slow apparent turnover of sedimentary IPL-iGDGT pools is consistent with substantial persistence and/or recycling of archaeal lipids and suggests that archaeal lipids may persist for extended periods after biosynthesis. These molecular biosignatures are highly resistant to degradation and can be well-preserved in terrestrial hydrothermal mineral matrices (Pancost et al., 2005; Kaur et al., 2011, 2015; Teece et al., 2020), although long-term preservation of iGDGTs in sinter remains to be demonstrated. Preservation may be further enhanced by rapid mineral entombment and protection within silica- or iron-rich deposits (Weimann et al., 2025). These observations support archaeal lipids as potentially durable biosignatures in hydrothermal environments on Mars. Although current rover-based instrumentation is unlikely to detect intact iGDGTs directly because of analytical limitations (Mahaffy et al., 2012; Beegle et al., 2015), targeting diagnostic cleavage products such as biphytane chains may improve the detectability of archaeal lipid biosignatures during *in situ* analyses. The detectability of these compounds on Mars remains speculative given current instrumental sensitivity and sample-processing constraints.

### 4.6 Conclusions

LH-SIP assays indicate that sedimentary IPL-iGDGT-derived biphytanes exhibit apparent turnover on decadal timescales in hot spring sediments. This slow apparent turnover likely integrates processes beyond de novo lipid synthesis, including persistence of older IPL-iGDGTs and recycling or re-functionalization of relict lipid components. These processes can decouple lipid turnover from cellular growth, such that apparent generation times likely overestimate true archaeal cell division times. Evidence for ongoing IPL-iGDGT production under near-natural incubation conditions, coupled with their chemical robustness, reinforces their value as persistent lipid biosignatures in hot spring sediments. Our findings provide the first empirical constraints on archaeal lipid production in hydrothermal spring sediments and offer a framework for interpreting iGDGT signatures in ancient terrestrial hydrothermal deposits with relevance to Mars exploration. Additional work is needed to determine how lipid recycling, persistence, and biosynthetic remodeling shape apparent iGDGT turnover in hydrothermal archaeal communities.

## Acknowledgments

This work was supported by the New Hampshire Space Grant Consortium (NHSGC) by NASA (to Dartmouth College, supporting CMH); Simons Foundation Award #623881 (WDL); NSF-EAR #1928303940 (WDL, SHK); Walter and Constance Burke Award (WDL); the Lewis and Clark Fund for Exploration and Field Research in Astrobiology, awarded by the NASA Astrobiology Program and the American Philosophical Society (CMH); and an Early Career Collaboration Award from the NASA Astrobiology Program (CMH). Yellowstone samples were collected under the National Park Service permit YELL-SCI-5544 (ESB). El Tatio samples were collected with permission from the Caspana and Toconce communities. We thank Dan Colman (MSU), Amanda Calhoun (Harvard) and Jenny Blamey (Biociencia Fundación Científica y Cultural, Chile) for their field support. We thank the Harvard Lab for Molecular and Biogeochemistry and Organic Geochemistry and the CU Boulder Earth Systems Stable Isotope Lab (CUBES-SIL) Core Facility (RRID:SCR_019300) for analytical contributions; and Susan Carter (Harvard), Amanda Calhoun (Harvard), and Ashley Maloney (CU Boulder) for expert laboratory support. This manuscript benefited from conversations with Tristan Caro (Caltech).

## Open Research Statement

All data and code pertaining to this manuscript are available are archived on Zenodo as *Hot Spring Lipid Hydrogen Stable Isotope Probing Analysis* (Version 1.0.1; Harris, 2026), https://doi.org/10.5281/zenodo.227C3239. The repository includes cleaned input datasets for GDGT and BP lipid abundances, BP hydrogen isotope composition, and experimental metadata, as well as all R scripts used for isotope label uptake and growth-rate calculations, uncertainty propagation, sensitivity analyses, and figure and table generation.

