## Supplemental Information for "Lipid Hydrogen Stable Isotope Probing Reveals Archaeal Lipid Turnover in Hot Spring Sediments"

*JGR Biogeosciences*

**Contents of this file**

Text S1 to S3

Figures S1 to S6

Tables S1 to S3

### Introduction

Included here are detailed derivations for uncertainty propagation in apparent growth-rate calculations, numerical sensitivity analyses that constrain the detection limits of the LH-SIP approach, and calculations used to estimate annualized production rates and new lipid production during the incubation. Supplementary tables provide full experimental summary data, literature compilations of archaeal lipid turnover estimates across environments, and literature compilations of laboratory growth rates determined for various archaeons isolated from hydrothermal environments.

### Supplementary Text

#### S1 Propagating error in growth rate and apparent generation time calculations

Archaeal lipid growth rates (μ, day$^{-1}$) and corresponding generation times ($T_G$, years) were calculated from $^2H_2O$ SIP experiments using a modified first-order exponential incorporation model (Caro et al., 2023) (see main text for additional details).

To calculate the uncertainty in $\mu$ (denoted $\sigma_\mu$), we propagated errors from all input parameters using a combination of partial derivatives and standard propagation rules. The final expression incorporates the analytical uncertainty of isotopic measurements ($F_0$, $F_t$, and $F_L$ for lipid compounds at time 0 and time t and the $^2H_2O$ labeling solution, respectively) and variability in assimilation efficiency (α):

Eq. (S1)
$$\sigma_\mu = \sqrt{\frac{\sigma_{F_0}^2 \cdot (\alpha F_L - F_t)^2 + \sigma_{F_T}^2 \cdot (F_0 - \alpha F_L)^2 + (F_0 - F_t)^2 \cdot (F_L^2 \cdot \sigma_\alpha^2 + \alpha^2 + \sigma_{F_L}^2)}{t^2 \cdot (F_0 - \alpha F_L)^2 \cdot (\alpha F_L - F_t)^2}}$$

All terms in the numerator account for contributions from each parameter's uncertainty.

- $\sigma_{F0}$, $\sigma_{Ft}$, and $\sigma_L$ : analytical precision of $^2H$ measurements
- $\sigma_\alpha$ : uncertainty in the assimilation efficiency, estimated from literature values

The resulting $\mu$ and $\sigma_\mu$ were converted to year$^{-1}$. Generation time uncertainty ($\sigma_{TG}$) was propagated using the derivative of the inverse function:

Eq. (S2)
$$\sigma_{T_G} = \frac{ln\,(2)}{\mu^2} \cdot \sigma_\mu$$

These calculations were applied to individual biological replicates for each compound (individual BP) and time point (3 or 14 days). Growth rates and generation times were then integrated over both time points. Propagated standard errors were calculated using weighted variances that included both measurement error and variability among biological replicates. This approach allowed us to quantify apparent lipid biosynthesis rates and their uncertainty under conditions of very low $^2H$ incorporation.

**S2 Determining the sensitivity and detection limits of LH-SIP incubations**

***Detection Limits*** We estimated the minimum detectable enrichment of $^2$H in lipids (i.e., the minimum $\Delta^2$H) that would yield a statistically resolvable growth rate under our experimental conditions. We calculated the propagated uncertainty in growth rate ($\sigma_\mu$) over a range of $\Delta^2$H values and defined the detection limit as the minimum value of $\Delta^2$H for which the estimated growth rate exceeded twice its propagated uncertainty (i.e., $\mu > 2\sigma_\mu$), consistent with a 95% confidence threshold. We considered a range of candidate $\Delta^2$H values generated from $F_t$ values between $F_0$ (i.e., no $^2$H-uptake) to $\alpha F_L$ (i.e., the theoretical maximum achievable $^2$H-uptake in newly synthesized lipids). Results reveal that lipids must be enriched by at least 1.5 ppm $^2$H (equivalent to 9.6 ‰, VSMOW increase in $\delta^2$H) over the incubation period for the growth rate (μ) to be statistically distinguishable from zero at the 95% confidence level (Figure S5). Over a 14-day incubation, this corresponds to a minimum detectable growth rate of 0.017 ± 0.008 year$^{-1}$ and a maximum detectable apparent generation time of 41.8 ± 20.9 years.

***Sensitivity Analysis*** In general, stronger $^2H_2O$ labeling lowers the incubation time needed to detect a given lipid enrichment, and for a fixed incubation time, stronger labeling permits detection of longer apparent generation times. We performed a sensitivity analysis that models the incubation time needed to detect theoretical $^2$H enrichment in archaeal lipids over a range of labeling and physiological scenarios. We assumed exponential isotopic incorporation according to the following equation:

Eq. (S3)
$$\Delta^2 H = F_t - F_0 = (1 - e^{-r \cdot t}) \cdot (\alpha \cdot F_L - F_0)$$

Where *r* is the specific biosynthesis rate (1/days), *t* is the duration of the incubation (days); α is the assimilation efficiency and fractionation of water hydrogen during lipid biosynthesis; and $F_0$, $F_t$, and $F_L$ are the fractional abundances of $^2$H in biphytanes at time 0, biphytanes at time *t*, and in the isotopically labeled spring water, respectively. Assimilation efficiency, α, represents the fraction of iGDGT hydrogen sourced from water (as opposed to the organic substrate used for growth), which also contributes H to lipids during heterotrophic growth.

To determine the impact of $^2H_2O$ labeling strength on assay detection limits, we performed simulations with three labeling solutions: $F_L$ = 0.37 atom % $^2$H (the labeling strength achieved at Beryl), 1 atom % $^2$H, and 10 atom % $^2$H; labeling solutions below 15 atom % are recommended to avoid $^2H_2O$ toxicity (Berry et al., 2015; Kopf et al., 2016; Figure S6). In each simulation, we consider two boundary conditions for assimilation efficiency: α = 0.56 for pure heterotrophy, α = 0.76 for pure autotrophy. We consider incubation durations of 0.01

days (~15 minutes) to 100 days. We show lipid $^2H$ enrichments ($\Delta^2H$) of 1.56 to 1556 ppm (equivalent to $\delta^2H$ increases of 10 ‰ to 10,000 ‰ VSMOW), which encompasses minimum detectable $\Delta^2H$ and the analytical sensitivity of typical GC-HTC-IRMS methods. For each simulation, we calculated the incubation time necessary to detect growth by archaea with certain generation times.

Under the LH-SIP conditions used in this study, ($F_L$ = 0.37 atom % $^2H$, 3-14 day incubations; Figure S6A), the detectable generation time ranged from ~2 days to ~30 years (heterotrophic scenario) or ~2 days to ~40 years (autotrophic scenario). These results define the minimum and maximum archaeal generation times that can be resolved with the $^2H_2O$-SIP protocol used on hot spring sediments in this study and help constrain the limits of detection in low-activity environments. Future studies using more highly enriched labeling solutions (e.g., 1 to 10 atom % $^2H$) over 14-day incubations, could extend the range of detectable generation times to ~740 years (heterotrophic scenario) to ~1000 years (autotrophic scenario) (Figure S6B, C).

**S3 Estimating new lipid production and annual production rate**

To estimate the amount of new lipid biomass synthesized during LH-SIP incubations, we used exponential growth modeling based on the growth rates (μ) derived in Eq. S3. The total new biomass synthesized during the incubation was calculated for individual biphytane compounds and aggregated across all compounds using the following equation:

Eq. (S4) $$B_{new} = B_0 \cdot (e^{\mu t} - 1) \cdot 10^3$$

Where $B_{new}$ is new biphytane production over the incubation (ng $g^{-1}$ sediment), $B_0$ is the initial biphytane concentration (μg $g^{-1}$ sediment), μ is the growth rate returned our SIP incubations ($year^{-1}$), *t* is the incubation time in years. $B_t$ is converted to a % increase relative to $B_0$. This model assumes that lipid biosynthesis scales proportionally with biomass increase and proceeds at a constant specific rate over the incubation.

To account for uncertainty in growth rate, we propagated error using the derivative of the exponential function with respect to μ:

Eq. (S5) $$\sigma_{B_{new}} = |\, B_0 \cdot e^{\mu t} \cdot t| \cdot \sigma_\mu$$

$B_{new}$ is best interpreted as an estimate of potential new biomass production under constant exponential growth assumptions. These calculations represent potential new biphytane production under the simplifying assumption that isotope incorporation reflects *de novo* lipid synthesis proportional to biomass production. They do not account for persistence of older IPLs, recycling or re-functionalization of pre-existing lipid components, or biosynthetic lag. If these processes are important, the calculated values should be interpreted as apparent lipid production estimates rather than direct biomass production rates.

### Supplementary Figures

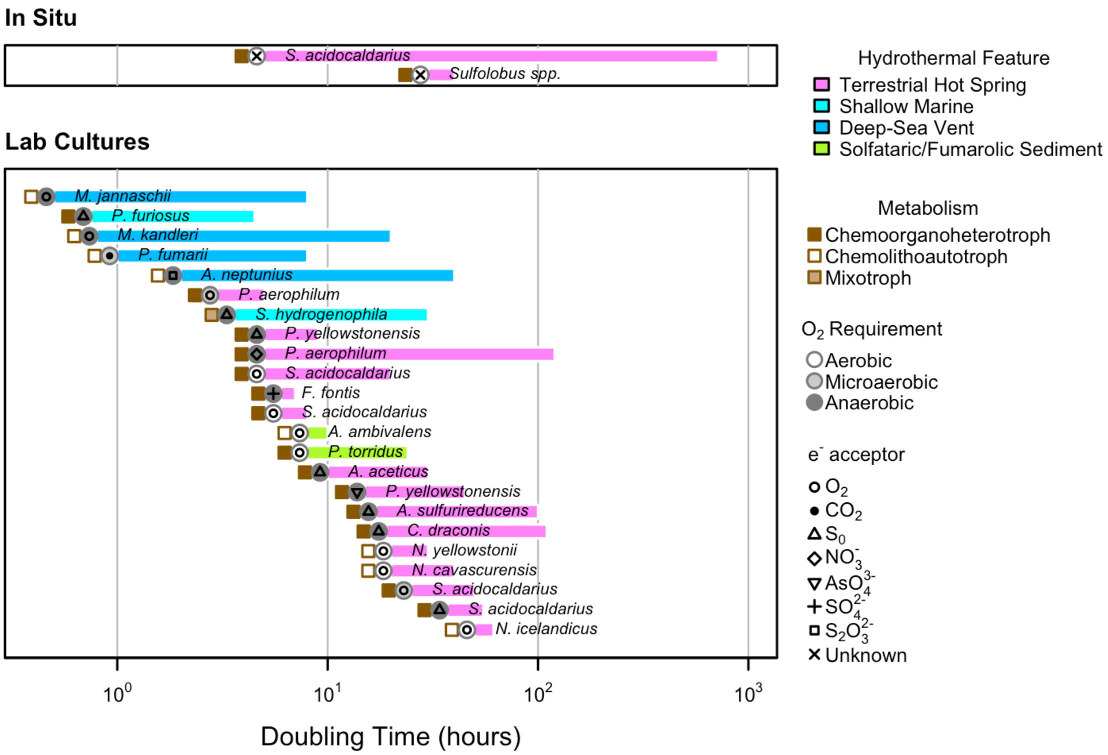


***Figure S1.***

Summary of archaeal doubling times determined via *in situ* measurements in hot springs (top panel) and for laboratory culture studies of isolates (bottom panel). Culture data includes archaeal strains isolated from various hydrothermally influenced environments indicated by bar color (including terrestrial hot spring, shallow marine hot brine seep, deep-sea vent, or solfataric/fumarolic sediment) cultured in controlled laboratory experiments. Symbols plotted at the left end of each bar denote, from left to right, metabolic mode, oxygen requirement, and terminal electron acceptor, as summarized in the legend. Lab cultures include measurements made under experimentally defined conditions that often varied temperature, pH, and/or substrate availability and typically approximate optimal or near-optimal growth conditions for the organism. Shown for comparison are the only two *in situ* archaeal growth-rate estimates identified for hydrothermal systems, both from acidic hot springs in Yellowstone National Park. These estimates were generated via tracer dilution (washout) rates and cell counts, or by measuring colonization on settlement plates. Across all compiled studies, reported doubling times span more than three orders of magnitude, from <1 hour to >500 hours, highlighting the broad physiological diversity of thermophilic and hyperthermophilic archaea across hydrothermal settings and metabolic strategies. Full metadata for each entry are provided in Table S3.

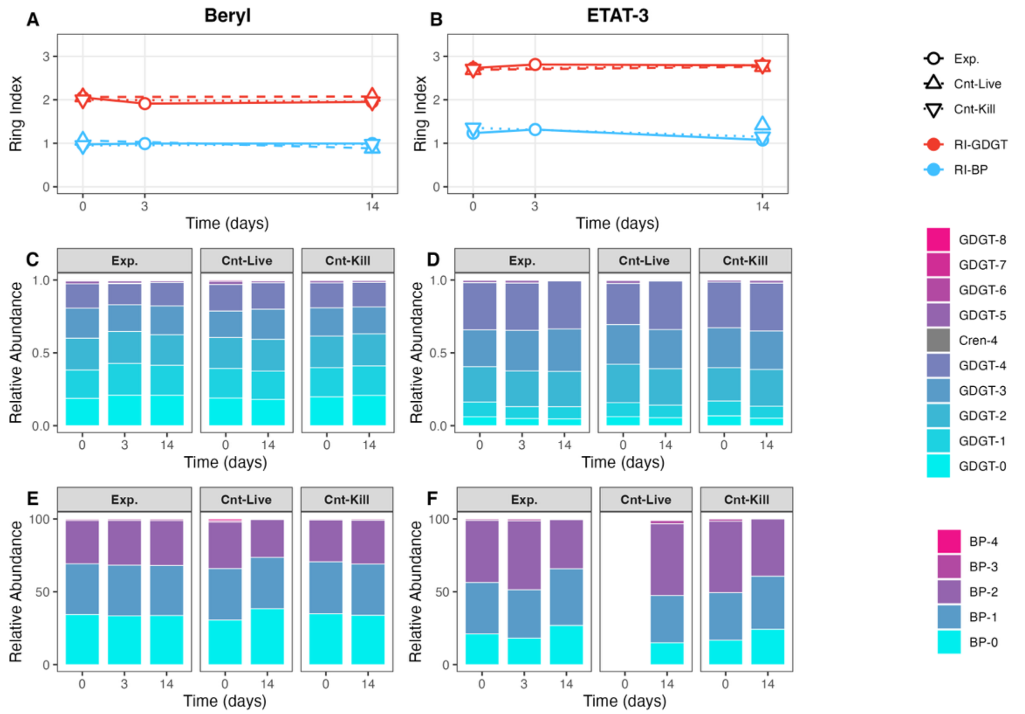


***Figure S2.***
IPL lipid distributions and Ring Indices over LH-SIP incubation time course for Beryl and 'ETAT-3' sediments. (A, B) Ring Index calculated for iGDGT lipids (red lines) and iGDGT-derived BPs (blue lines) for experimental replicates (circles) and controls (triangles) over the LH-SIP incubation. Points show means across biological duplicates; SE is smaller than symbols in all cases. (Distribution of iGDGT lipids (C, D) and BP lipids (E, F) for experimental replicates and controls. No major shifts in lipid composition occurred during the incubation at either site.

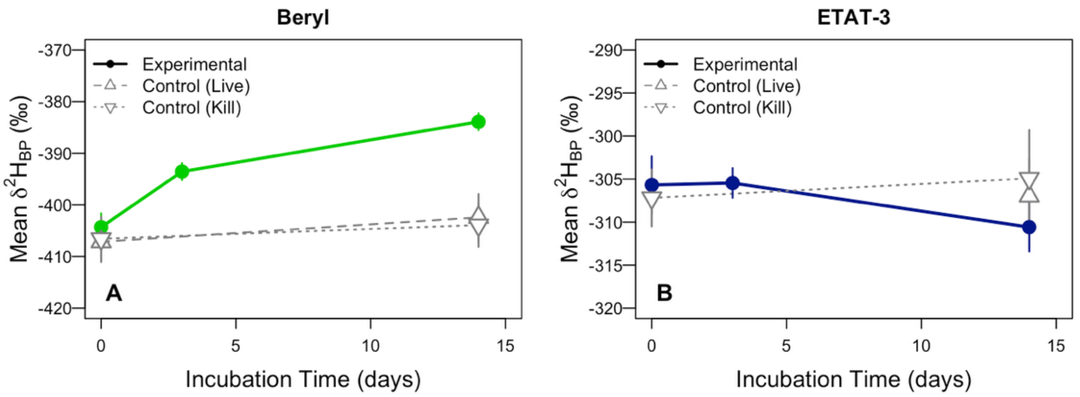


***Figure S3.***

Mean $\delta^2H_{BP}$ composition (‰) of archaeal lipids over the LH-SIP incubation time course for experimental and control samples averaged across biphytane moieties for Beryl (A) and 'ETAT-3' (B) sediments. After 14-day incubations, archaeal IPL lipids isolated from Beryl sediments were enriched by 20.4 ± 3 ‰ relative to the starting composition at $T_0$. No enrichment was observed at 'ETAT-3' over the incubation.

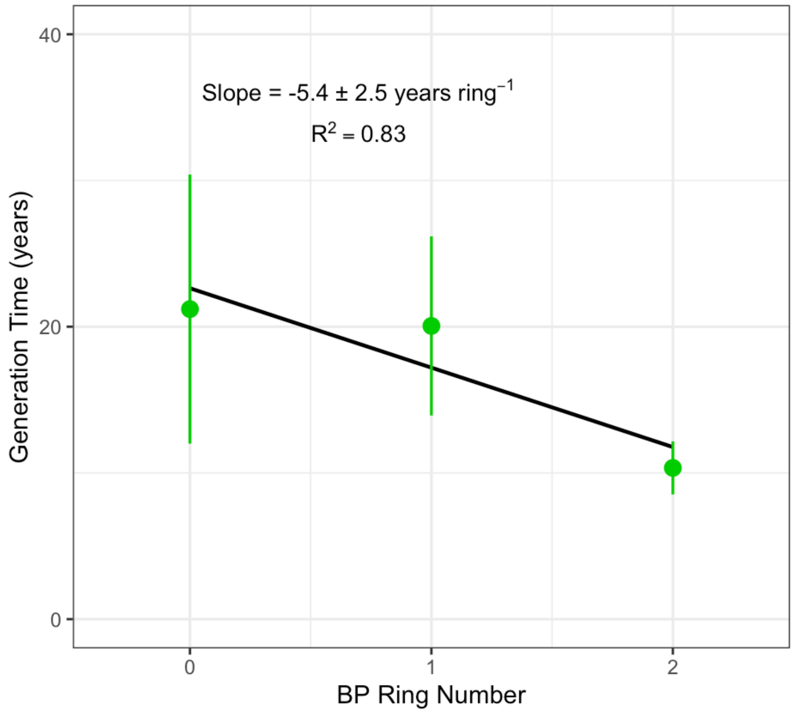


***Figure S4.***

Linear regression of biphytane ring number versus apparent generation time. Points and error bars show mean values and propagated uncertainty from isotopic measurements, tracer assimilation efficiencies, and biological replication. The shaded region shows the 95% confidence interval of the regression. On average, the addition of each cyclopentane ring decreases the apparent generation time by 5.4 ± 2.5 years. This trend suggests that more highly cyclized biphytanes may be associated with more actively synthesized or more rapidly cycling archaeal lipid pools in hot spring sediments.

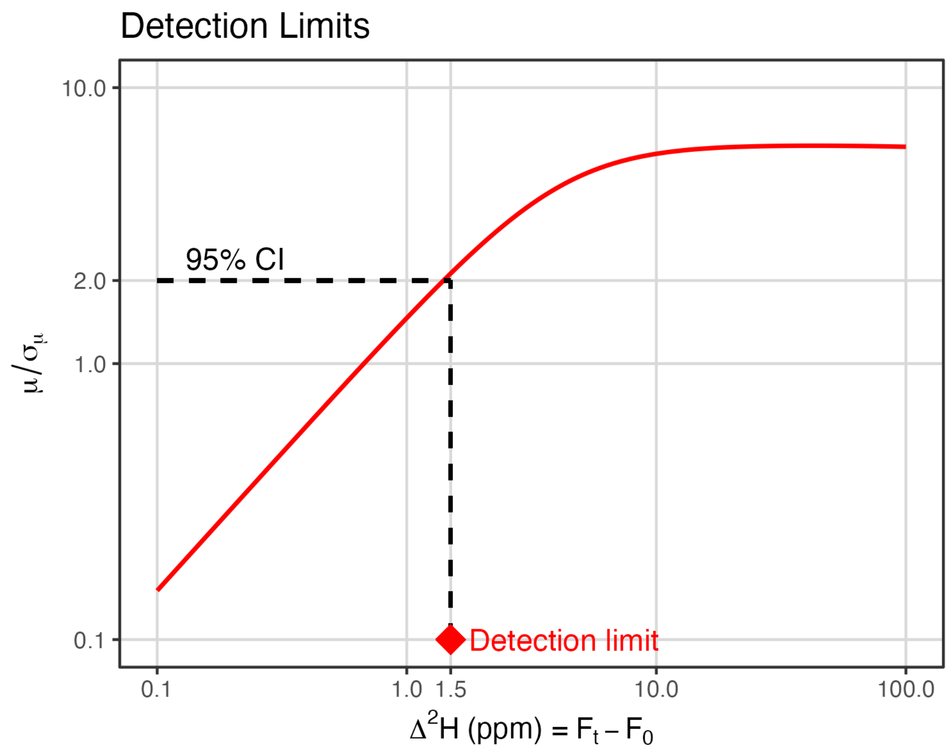


***Figure S5.***

Theoretical lipid $^2$H enrichment (ppm) vs. corresponding ratio of growth rate to propagated uncertainty ($\mu / \sigma_\mu$) given the parameters and uncertainties in LH-SIP assays performed in this study. The dashed lines indicate the detection limit (defined as the 2σ threshold for statistically significant detection of $^2$H-uptake into lipids), which is achieved with ≥ 1.5 ppm $^2$H enrichment relative to $t_0$, (equivalent to enrichment of ≥ 9.6 ‰, VSMOW).

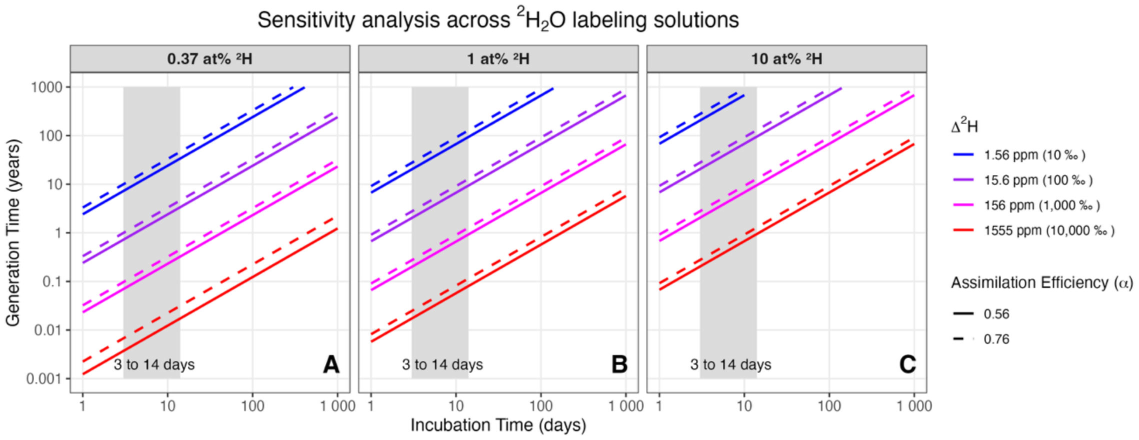


***Figure S6.***
Sensitivity analysis to determine the incubation time necessary to detect growth by microbes with apparent generation times ranging from hours to 1000 years in LH-SIP assays. We consider $^2H_2O$ labeling solutions with three $F_L$ values: (A) 0.37 atom % $^2H$ (3,700 ppm $^2F$), which was the label strength achieved in this study; (B) 1 atom % $^2H$ (10,000 ppm $^2F$); and (C) 10 atom % $^2H$ (100,000 ppm $^2F$). Colored lines denote a plausible range of lipid $^2H$-enrichments ($\Delta^2H$ values) from +10 ‰ (~1.6 ppm, the assay detection limit) to +10,000 ‰ (~1555 ppm). The line type indicates the boundary conditions for assimilation efficiency (α): pure heterotrophy (α = 0.56; solid lines) and pure autotrophy (α = 0.76; dashed lines). Given the 3 to 14-day incubations and LH-SIP parameters used in this study (A), we can detect apparent generation times of < 1 days to 42 years. Increasing the strength of the $^2H_2O$ labeling solution to 1 atom % $^2H$ (B) would allow detection of generation times from 6 days to 91 years. Increasing to a 10 atom % $^2H$ solution (C) would allow detection of generation times from 74 days to 1000 years.

### *Supplementary Tables*

***Table S1.***

Summary data for *ex situ* LH-SIP experiments on hot spring sediments including live and killed controls and experimental replicates for each time point. The relative abundance (%) of IPL-iGDGT-derived biphytane moieties and BP Ring Index is shown. The lipid $\delta^2$H composition (‰) for each BP moiety, the BP abundance-weighted mean $\delta^2$H, and the mean ring difference ($\Delta\delta$/ring, ‰) is also shown.

| | | | | Relative Abundance (%) | | | | | | Lipid $\delta^2$H (‰) | | | | $\Delta\delta^2$H/ring (‰) |
|---|---|---|---|---|---|---|---|---|---|---|---|---|---|---|
| Site | Type | Time Point (days) | N | BP-0 | BP-1 | BP-2 | BP-3 | BP-4 | Ring Index | BP-0 | BP-1 | BP-2 | Wt. Mean | Mean |
| Beryl | Killed Control | 0 | 1 | 35 | 36 | 29 | 1 | 0 | 0.95 | -409 ± 9.3 | -413 ± 6.6 | -396 ± 5 | -407 ± 4.3 | 8 ± 6.8 |
| | | 14 | 1 | 34 | 35 | 30 | 1 | 0 | 0.98 | -409 ± 9.3 | -411 ± 6.6 | -389 ± 5 | -404 ± 4.2 | 12 ± 6.8 |
| | Live Control | 0 | 1 | 31 | 35 | 32 | 1 | 1 | 1.07 | -413 ± 9.3 | -414 ± 5.8 | -393 ± 5 | -407 ± 3.9 | 12 ± 6.6 |
| | | 14 | 1 | 38 | 35 | 26 | 0 | 0 | 0.88 | -406 ± 9.3 | -409 ± 6.6 | -388 ± 5.5 | -402 ± 4.5 | 11 ± 6.9 |
| | Experimental | 0 | 2 | 34 ± 0 | 35 ± 0 | 30 ± 0 | 1 ± 0 | 0 ± 0 | 0.97 ± 0.01 | -412 ± 5.2 | -409 ± 2.6 | -390 ± 6.8 | -404 ± 4.8 | 13 ± 1 |
| | | 3 | 2 | 33 ± 1 | 35 ± 1 | 31 ± 1 | 1 ± 0 | 0 ± 0 | 0.99 ± 0.03 | -403 ± 1.9 | -401 ± 1.4 | -374 ± 0 | -394 ± 1.5 | 17 ± 1.1 |
| | | 14 | 2 | 34 ± 0 | 34 ± 1 | 31 ± 1 | 1 ± 0 | 0 ± 0 | 0.99 ± 0.01 | -397 ± 1.6 | -392 ± 3 | -360 ± 3.4 | -384 ± 3 | 22 ± 1.1 |
| 'ETAT-3' | Killed Control | 0 | 1 | 17 | 33 | 49 | 1 | 0 | 1.35 | -320 ± 8.2 | -308 ± 5.8 | -302 ± 4.6 | -307 ± 3.3 | 9 ± 5.6 |
| | | 14 | 1 | 24 | 36 | 39 | 0 | 0 | 1.15 | -311 ± 15.4 | -306 ± 8.7 | -301 ± 7 | -305 ± 5.6 | 6 ± 9.6 |
| | Live Control | 0 | 1 | - | - | - | - | - | - | - | - | - | - | - |
| | | 14 | 1 | 15 | 32 | 49 | 2 | 1 | 1.42 | -321 ± 15.4 | -301 ± 6.6 | -307 ± 5.5 | -307 ± 4.3 | 7 ± 8.9 |
| | Experimental | 0 | 2 | 21 ± 1 | 35 ± 0 | 43 ± 2 | 1 ± 0 | 0 ± 0 | 1.23 ± 0.03 | -314 ± 6.9 | -306 ± 3.3 | -301 ± 1 | -306 ± 3.3 | 7 ± 3.2 |
| | | 3 | 2 | 18 ± 3 | 33 ± 2 | 47 ± 4 | 1 ± 1 | 0 ± 0 | 1.32 ± 0.08 | -314 ± 5.5 | -307 ± 4.7 | -301 ± 4 | -305 ± 4.9 | 7 ± 0.8 |
| | | 14 | 2 | 27 | 39 | 34 | 0 | 0 | 1.08 | -322 ± 5.8 | -310 ± 4.6 | -303 ± 4.4 | -311 ± 2.8 | 10 ± 4.4 |

***Table S2.***

Comparison of archaeal IPL lipid turnover and generation times across diverse environments calculated from stable isotope probing (SIP), radiocarbon dating, modeling, and tracer experiments (data summarized in Figure 5). Methodologies include $^{13}$C-, $^{2}$H-, and $^{14}$C-based labeling or decay, as well as kinetic and box modeling. The table includes both *in situ* and *ex situ* estimates of archaeal lipid turnover in terrestrial soils, ocean water column, and marine surface sediments and deep subsurface. Values represent either estimated generation times or degradation half-lives (converted to years), with bounds indicating the reported range of values for each study. This study provides the first LH-SIP constraints on apparent turnover of archaeal IPL-iGDGT-derived biphytanes in terrestrial hydrothermal sediments. GlcDGD = Glycosyl diphytanylglycerol diether.

| Environment | Sample Description | Compound | Turnover / Generation Time | Method | Reference |
|---|---|---|---|---|---|
| Hot Spring Sediments | High temperature, circumneutral springs | BP-0 to -2 | 8-20 years, >42 years | $^{2}H_2O$ SIP | ***This study*** |
| Ocean | North Sea (Wadden Sea) | iGDGT-1 & Crenarchaeol | 76 to 128 d | $^{13}$C-DIC SIP | Wuchter et al., (2003); Pitcher et al., (2011) |
| Temperate soils | Forested mineral soils in Swiss alps | Pooled iGDGT | 1,400 to 2,000 years (10-20 cm); 2,000 to 6,000 years (50+ cm) | $^{14}$C dating of pooled iGDGTs | Gies et al., (2021) |
| Marine sediments | Methane rich, hydrothermally heated sediment from Guaymas Basin | Archaeol & iGDGTs | 380 days for archaeols to 28 years for various iGDGT compounds | $^{2}H_2O$ SIP | Kellermann et al., (2016) |
| | Marine shelf sediment, Iceland | iGDGT-0 & Crenarchaeol | ≥ 4 to 9 years | $^{13}$C-DIC SIP | Lengger et al., (2014) |
| | Deep Sea sediments, Sagami Bay, Japan | Caldarchaeol & Crenarchaeol | $^{13}$C incorporation into glycerol backbone after 9 days; No $^{13}$C labeling in isoprenoid chains (biphytanes) after 400 days | $^{13}$C-glucose SIP | Takano et al., (2010) |
| | Continental margin sediments (model to 1 km depth) | GlcDGD | Half-life up to $3.1\times10^5$ years for archaeal IPLs at depth; community generation time 1.6 - 73 kyr (below 1 m) | $^{14}$C-radiolabeled lipid decay assay + kinetic modeling | Xie et al., (2013) |
| | Surface sediment from the Peru Margin; subsurface sediments from Hydrate Ridge, Juan de Fuca Ridge | iGDGT-0 to -5 | Degradation of IPLs with a half-life increasing from 1 kyr at the sediment surface to 500 kyr at 1000 m | Model of IPL-iGDGT sediment concentrations | Lipp and Hinrichs (2009) |
| Marine subsurface | Anoxic sediments, Fjord Himmersfj ärden (Sweden) | BP-0 to -3 | Community turnover of 160 to 840 years | Dual $^{2}H_2O$ + $^{13}$C-DIC SIP | Wegener et al., (2012) |
| | Marine subsurface sediment (~10 m below seafloor) | 2G-IPLs iGDGT-0 to -2; Diglycosyl GDGTs | 1,700 - 20,500 years | $^{13}$C-substrate SIP | Lin et al., (2013) |

***Table S3.***

Source data and metadata for archaeal doubling times compiled in Figure S1. Entries include hydrothermal setting, species, metabolic mode, metabolism (including terminal electron acceptor where reported), oxygen requirement, doubling-time range (hours), measurement method, isolate collection context, and literature citation.

| Hydrothermal Setting | Species | Metabolic Mode | Metabolism | Oxygen Requirement | Doubling Time (hours) | Method | Collection Details | Reference |
|---|---|---|---|---|---|---|---|---|
| Terrestrial Hot Spring | *Sulfolobus spp.* | Chemoorganoheterotroph | Unknown | Aerobic | 30 to 40 | *In situ* | Yellowstone (USA) – 4 springs, Temp 55-92°C, pH 0.9 to 3.0; moving and still water | Bohlool and Brock, 1974 |
| Terrestrial Hot Spring | *S. acidocaldarius* | Chemoorganoheterotroph | Unknown | Aerobic | 5 to 720 | *In situ* | Yellowstone (USA) – small acidic spring (pH ~2.5, ~72 °C, sulfur-rich) | Mosser et al., 1974 |
| Deep-Sea Vent | *Methanocaldococcus jannaschii* | Chemolithoautotroph | $H_2 + CO_2$ | Anaerobic | 0.5 to 8 | Lab culture | Deep-sea hydrothermal vent (East Pacific Rise) | Susanti et al., 2019 |
| Deep-Sea Vent | *Methanopyrus kandleri* | Chemolithoautotroph | $H_2 + CO_2$ | Anaerobic | 0.8 to 20 | Lab culture | Deep-sea hydrothermal vent sediment | Huber, 1989 |
| Deep-Sea Vent | *Archaeoglobus neptunius* | Chemolithoautotroph | $H_2 + S_2O_3^{2-}$ | Anaerobic | 2 to 40 | Lab culture | Mid Atlantic Ridge - deep-sea vent | Slobodkina, 2021 |
| Deep-Sea Vent | *Pyrolobus fumarii* | Chemolithoautotroph | $H_2 + NO_3^-$ | Microaerobic | 1 to 8 | Lab culture | Mid Atlantic Ridge - Guaymas basin, black smoker | Anderson et al., 2011; Blöchl et al., 1997 |
| Shallow Marine | *Pyrococcus furiosus* | Chemoorganoheterotroph | Organics + $S^0$ | Anaerobic | 0.75 to 4.5 | Lab culture | Vulcano Island (Italy) - various beaches | Fiala and Stetter, 1989 |

| | | | | | | | | |
|---|---|---|---|---|---|---|---|---|
| Shallow Marine | *Stetteria hydrogenophila* | Mixotroph | $H_2$ + peptides / $S^0$ | Anaerobic | 3.6 to 30 | Lab culture | Paleohori Bay (Greece) - hot brine seep | Jochimsen, 1997 |
| Solfataric/Fumarolic Sediment | *Acidianusambivalens* | Chemolithoautotroph | $S_4O_6^{2-}$ + $O_2$ | Aerobic | 8 to 10 | Lab culture | Iceland - Leirhnjúkur hot spring, solfataric mud | Protze et al., 2011 |
| Solfataric/Fumarolic Sediment | *Picrophilus torridus* | Chemoorganoheterotroph | Organics + $O_2$ | Aerobic | 8 to 24 | Lab culture | Owakudani (Japan) - Dry solfataric soil | Schleper et al., 1995 |
| Terrestrial Hot Spring | *Nitrosocaldus yellowstonii* | Chemolithoautotroph | $NH_3$ + $O_2$ | Aerobic | 20 to 30 | Lab culture | Yellowstone (USA) - various springs | De La Torre et al., 2008 |
| Terrestrial Hot Spring | *Nitrosocaldus cavascurensis* | Chemolithoautotroph | $NH_3$ + $O_2$ | Aerobic | 20 to 40 | Lab culture | Italy - Ischia hot spring | Abby et al., 2018 |
| Terrestrial Hot Spring | *Nitrosocaldus icelandicus* | Chemolithoautotroph | $NH_3$ + $O_2$ | Aerobic | 49.92 to 61.44 | Lab culture | Iceland – Graendalur valley geothermal spring (6.5 and a temperature of 73°C) | Daebeler et al., 2018 |
| Terrestrial Hot Spring | *Sulfolobus acidocaldarius* | Chemoorganoheterotroph | Organics + $O_2$ | Aerobic | 5 to 20 | Lab culture | Yellowstone (USA) – acidic spring (pH 2–3, ~75 °C); lab culture in rich medium | Harris et al., 2025 |
| Terrestrial Hot Spring | *Sulfolobus acidocaldarius* | Chemoorganoheterotroph | Organics + $O_2$ | Aerobic | 6 to 8 | Lab culture | Yellowstone (USA) – acidic spring (pH 2–3, ~75 °C); lab culture in rich medium | Brock et al., 1972 |
| Terrestrial Hot Spring | *Acidilobus aceticus* | Chemoorganoheterotroph | Organics + $S^0$ | Anaerobic | 10 to 30 | Lab culture | Kamchatka (Russia) – acidic spring (pH ~3.5, ~85 °C) | Prokofeva et al., 2014 |
| Terrestrial Hot Spring | *Pyrobaculum yellowstonensis* | Chemoorganoheterotroph | Organics + $AsO_4^{3-}$ | Anaerobic | 15 to 45 | Lab culture | Yellowstone (USA) – Joseph's Coat Hot Spring (circumneutral, 80 °C, pH 6.1; arsenic ~0.135 mM) | Jay et al., 2015 |
| Terrestrial Hot Spring | *Acidilobus sulfurireducens* | Chemoorganoheterotroph | Organics + $S^0$ | Anaerobic | 17 to 100 | Lab culture | Yellowstone (USA) - Dragon Spring, cultures at pH 3, T 65 to 80°C | Boyd et al., 2007 |

| | | | | | | | | |
|---|---|---|---|---|---|---|---|---|
| Terrestrial Hot Spring | *Caldisphaera draconis* | Chemoorganoheterotroph | Organics + $S^0$ | Anaerobic | 19 to 110 | Lab culture | Yellowstone (USA) - Dragon Spring, cultures at pH 3, T 65 to 80°C | Boyd et al., 2007 |
| Terrestrial Hot Spring | *Sulfolobus acidocaldarius* | Chemoorganoheterotroph | Organics + $S^0$ | Anaerobic | 37 to 55 | Lab culture | Yellowstone (USA) – acidic spring (pH 2–3, ~75 °C); lab culture on sulfur only | Shivers & Brock, 1973 |
| Terrestrial Hot Spring | *Pyrobaculum aerophilum* | Chemoorganoheterotroph | Organics + $NO_3^-$ | Anaerobic | 5 to 120 | Lab culture | Iceland – hyperthermal spring (optimum 100 °C, vent water) | Völkl et al., 1993 |
| Terrestrial Hot Spring | *Pyrobaculum yellowstonensis* | Chemoorganoheterotroph | Organics + $S^0$ | Anaerobic | 5 to 9 | Lab culture | Yellowstone (USA) – Joseph's Coat Hot Spring (circumneutral, 80 °C, pH 6.1; arsenic ~0.135 mM) | Jay et al., 2015 |
| Terrestrial Hot Spring | *Fervidicoccus fontis* | Chemoorganoheterotroph | Organics + $SO_4^{2-}$ | Anaerobic | 6 to 7 | Lab culture | Kamchatka (Russia) – Uzon Caldera hot spring (slightly acidic, 65–70 °C) | Perevalova et al., 2010 |
| Terrestrial Hot Spring | *Sulfolobus acidocaldarius* | Chemoorganoheterotroph | Organics + $O_2$ | Microaerobic | 25 to 50 | Lab culture | Yellowstone (USA) – acidic spring (pH 2–3, ~75 °C); lab culture in rich medium | Harris et al., 2025 |
| Terrestrial Hot Spring | *Pyrobaculum aerophilum* | Chemoorganoheterotroph | Organics + $O_2$ | Microaerobic | 3 to 5 | Lab culture | Iceland – hyperthermal spring (optimum 100 °C, vent water) | Völkl et al., 1993 |

/end
